# Design and Application of a µSiM Outer Blood-Retinal Barrier (OBRB) Model as a Drug Development Tool

**DOI:** 10.64898/2026.08.23.746552

**Authors:** Kevin C. Ling, Jordan Jones, Gram Hepner, Ahmet Gurcan, Rufaro Gamariel, Andres Muriel-Torres, Meng-chun Hsu, Mehran Mansouri, Sami Farajollahi, Vinay Abhyankar, Ruchira Singh, Danielle Benoit, James McGrath

## Abstract

The outer blood-retinal barrier (OBRB) is the primary interface through which systemically circulating drugs reach the retina. A tool that measures delivery across this barrier would support the development of targeted therapies as alternatives to repeated intravitreal injection, and the screening of drugs that reach the retina as an off-target toxicity. Such a tool should deliver drugs fluidically through a vascular compartment, measure transport across the retinal pigment epithelium (RPE), and display disease phenotypes relevant to efficacy. Here we adapt the µSiM platform, which places epithelium and endothelium in direct juxtaposition across a permeable, optically transparent silicon nitride nanomembrane. ARPE-19 and human umbilical vein endothelial cells (HUVECs) were used as development cell sources. ARPE-19 monocultures reached a transepithelial electrical resistance of 68 ± 26 Ω cm^2^ by 28 days, and ARPE-19 + HUVEC co-cultures reached a small-molecule permeability of 6.34 ± 1.3 × 10^−4^ cm min^−1^ within 14 days, a state reported elsewhere only after longer culture. The barriers developed an intervening basement membrane. Drugs perfused through the basal vascular channel crossed into an open apical well, where sampling and mass spectrometry showed transport correlating with lipophilicity, as reported in vivo. The device also displayed two clinically relevant phenotypes. Digoxin at a clinically toxic concentration reduced viability in the co-barrier by about half and doubled permeability. In a vascularized configuration, VEGF drove endothelial invasion of the RPE layer, as seen in neovascular AMD. The µSiM-OBRB therefore satisfies basic design criteria for measurement of drug bioavailability, toxicity, and efficacy.

## Introduction

The outer blood-retinal barrier (OBRB) is the primary interface that controls which systemically circulating drugs reach the retina. It is a three-layered structure: a fenestrated vasculature (the choriocapillaris); an intervening basement membrane (Bruch’s membrane), synthesized by both the vascular endothelium and the retinal pigment epithelium (RPE); and the RPE itself, whose tight junctions and transporters make it the rate-limiting layer^1,2^ (**Fig. 1A**). A drug reaching the outer retina from the systemic circulation must leave the choriocapillaris, traverse Bruch’s membrane, and cross the RPE monolayer; the combined permeability of these layers sets the ceiling on retinal drug bioavailability.^3^ Reconstructing this layered interface, together with the vascular route by which drugs arrive, is a basic requirement for any drug development tool intended to evaluate retinal drug bioavailability.

**Fig. 1.**
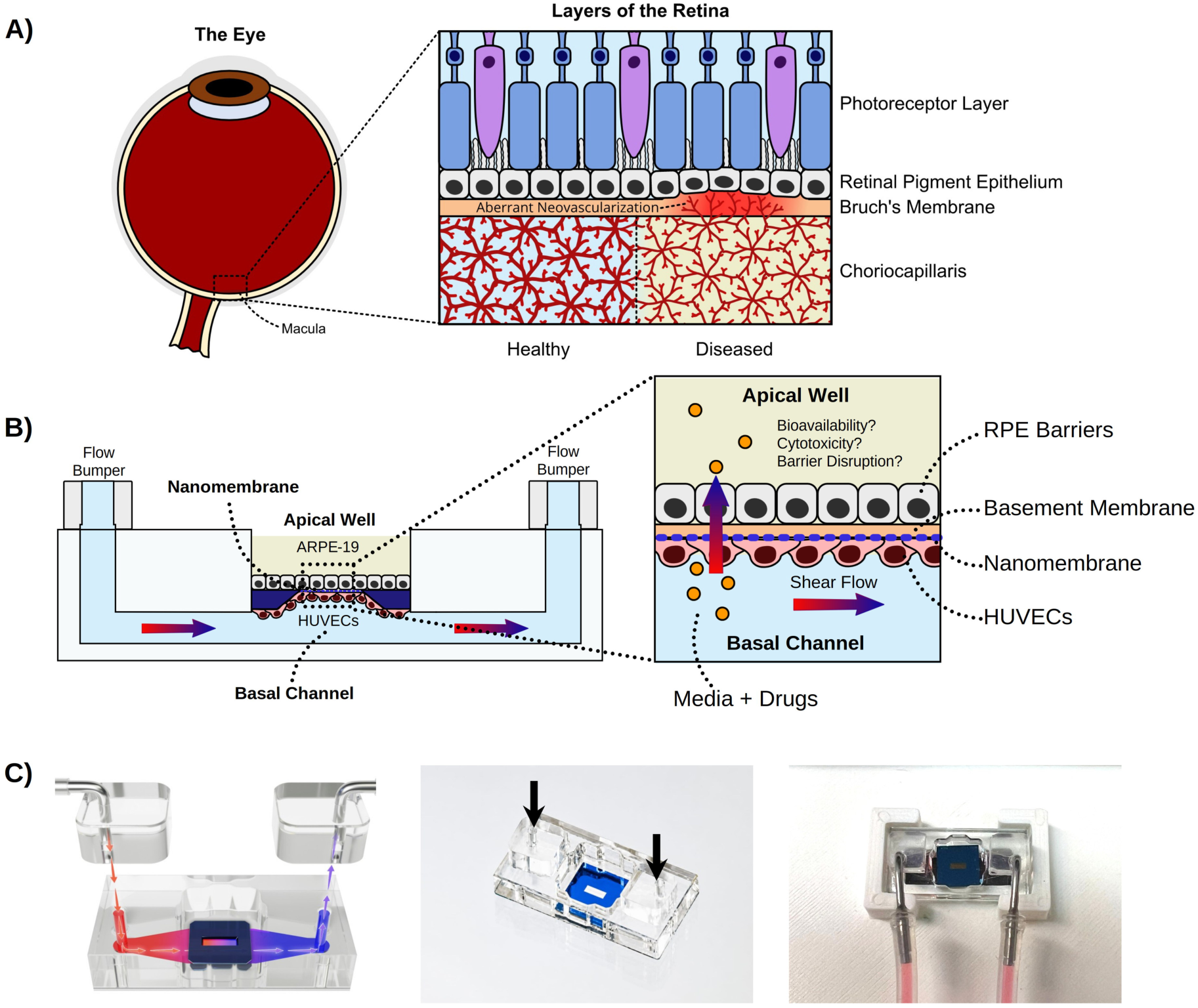
µSiM-OBRB enables in vitro modeling of retinal drug transport, drug toxicity, and pathological phenotypes of retinal degeneration. **(A)** Schematic of the outer blood- retinal barrier. Healthy and diseased states are shown; in the diseased retina, aberrant neovascularization from the choriocapillaris breaches Bruch’s membrane and invades the retinal pigment epithelium, a hallmark of neovascular AMD. **(B)** Diagram depicting ARPE-19 cells and HUVECs in a µSiM device. ARPE-19 form a cell barrier on top of a NPSN membrane. HUVECs are seeded in the bottom channel on the lower surface of the membrane, directly juxtaposed with apical ARPE-19 cells. ARPE-19-HUVEC co- barriers enable *in vitro* assessment of barrier function, retinal drug uptake, and toxicity in static and shear flow conditions. **(C)** Left; 3D render of *µSiM-OBRB* and two PDMS flow bumpers, with red fluid representing inward flow and blue fluid representing outward flow. Middle: photograph of µSiM-OBRB device with PDMS flow bumpers attached. Arrows denote placement of adhered PDMS flow bumpers. Right: photograph of µSiM-OBRB attached to a circulating fluidic circuit.

Numerous clinically approved drugs indicated for non-retinal conditions have been linked to retinal degeneration.^4–6^ They span cardiac glycosides, antimalarials, antiretrovirals, kinase inhibitors, and cytotoxic agents, as summarized in **Table S1†**. Hydroxychloroquine is the most consequential of them. It accumulates in the mammalian retina, binds melanin, and induces cell death in RPE cells,^7–13^ and as of 2019 roughly 167,000 patients were prescribed the drug, of whom more than 70,000 were long-term users at risk for retinal maculopathy.^14–16^ Although the mechanisms and time course of toxicity differ across these agents, and onset ranges from acute to chronic exposure, each must cross the inner or outer blood-retina barrier to reach the retina, either through permeation or direct damage. No clinically approved treatment can fully reverse toxicity-induced retinopathy, though stem-cell-based regenerative approaches are in clinical trials.^4,5,17–22^

Beyond off-target toxicity, the OBRB also determines whether drugs intended to treat the retina can reach it. Most approved retinal therapies, including the anti-VEGF biologics that are standard of care for neovascular age-related macular degeneration (AMD), are delivered by repeated intravitreal injection because systemically administered drugs cross the OBRB poorly.^23^ Intravitreal injection is invasive, carries risks such as endophthalmitis and retinal detachment, and burdens patients and clinics, motivating systemically administered, retina-targeted alternatives^24^ and in vitro tools for screening retinal pharmacokinetics. Pathological breakdown of the OBRB is itself a hallmark of neovascular AMD, in which choroidal vessels invade across the barrier and disrupt the RPE; AMD is both a leading cause of blindness and the leading indication for anti-VEGF therapy.^23^ A tool that reproduces this invasive phenotype could therefore evaluate not only barrier toxicity and bioavailability but also drug efficacy.

Current in vivo models are poorly suited for retinal drug screening. The retinae of mice differ structurally and functionally from those of humans: they lack a cone-rich macula, have thinner Bruch’s membrane, and their RPE cells have a higher relative phagocytic load.^25^ Non-human primates are physiologically closer but are impractical for mechanistic or therapeutic studies because of genetic inflexibility, slow disease progression, and high cost, and they still do not recapitulate all features of human retinal disease.^26–28^ Even human clinical trials are too short to capture the effects of chronic retinal drug exposure.^5,12^ As a result, both the mechanisms driving drug-induced retinal toxicity and the barrier behavior that governs retinal drug delivery remain poorly understood.

In vitro OBRB models capture parts of this challenge but not the whole. Static RPE permeability assays on porous filters quantify drug transport but lack a vascular compartment, flow, or an endothelium.^29–32^ Relative to other capillary networks and normalized for its size, the choriocapillaris is one of the most highly perfused microvascular networks in the human body,^33–38^ underscoring the importance of flow in models of the OBRB. Furthermore, flow disruptions in the choroid have also been linked to retinal degeneration.^39–41^ Microfluidic OBRB chips have added these elements: Arik et al. placed an immortalized RPE monolayer opposite a HUVEC microvessel across a porous polyester membrane and reported clinically relevant tissue-permeability and vascular-structure read-outs,^42^ and related microfluidic OBRB models drive wet-AMD-like choroidal neovascularization through hypoxic or VEGF stress.^29,43^ Organoid retina-on-chip platforms recapitulate drug toxicity, including chloroquine and gentamicin, and reveal an RPE protective effect,^44^ and 3D iPSC RPE–choriocapillaris constructs model AMD pathology and nominate disease factors.^45^ Each system achieves one or two of the functions a retinal drug development tool requires, and each separates RPE from endothelium with a comparatively thick porous membrane, a hydrogel, a matrix, or organoid architecture. However, none of these models place the epithelium and endothelium in direct juxtaposition or integrate quantitative microfluidic bioavailability measurement with toxicity and disease phenotypes in a single device. The tissue chip design presented here does both.

The modular **µSiM** (**micro**fluidic device featuring a **si**licon nitride **m**embrane) device platform was developed to enable use of 100 nm thick porous silicon nitride ‘nanomembranes’ in devices customized to fit a range of applications in diagnostics and tissue chips.^46^ For tissue chips, the µSiM has been applied to the development of a wide range of barrier MPS models including those for multiple sclerosis,^47,48^ sepsis,^49,50^ and tendon fibrosis.^51^ In each of these, the nanomembrane is used to create a model interface between a vascular compartment and a tissue compartment. The thinness and optical transparency of nanomembranes enables high-content, high-fidelity imaging. In addition to devices and protocols for establishing tissue models, we have developed a companion set of tools and methods for measuring barrier function and characterizing barrier and adjacent tissue through live imaging, immunofluorescence, and transcriptomics.^50,51^ The µSiM is modular by design: reconfiguring its components and exchanging the membrane adapt the same device into fit-for-purpose models of different tissue interfaces. Here, we employ the µSiM to model the outer blood-retina barrier (**µSiM-OBRB**) (**Fig. 1B, C**). The nanomembrane forms the epithelial-endothelial interface, continuous perfusion of the basal channel delivers drug to the vascular side, and an open apical well exposes the retinal side for sampling.

We deliberately limit the scope of this work to the design and basic operation of the device and reserve the final cell composition for the next stages of development. Specifically, we use ARPE-19 and human umbilical vein endothelial cells (HUVECs) as convenient, reliable development cell sources, recognizing that sourcing the most physiologically faithful cells, iPSC-derived RPE and tissue-specific or iPSC-derived choroidal endothelium, is a challenge reserved for biological validation in a next-generation model.

Here, we validate device function, which requires only basic cell performance against three engineering objectives: the device must (1) deliver a drug panel through the perfused vascular channel and quantify its transport to the retinal compartment (bioavailability), (2) detect drug-induced loss of barrier integrity (barrier toxicity), and (3) reproduce a neovascular disease phenotype (neovascularization). To this end, we show that ARPE-19 cultured in µSiM-OBRB formed tight, polarized barriers, and that ARPE- 19 + HUVEC co-barriers reach mature permeability within two weeks. Mass spectrometry was used to assess the pharmacokinetics of a panel of small-molecule drugs through the basal vascular channel, across the RPE-endothelial barriers, and into the apical retinal chamber, recapitulating lipophilicity-dependent drug transport across the native OBRB. Digoxin induced a barrier-disruption phenotype. By switching to a microporous membrane and integrating a fibrin hydrogel-entrapped 3D microvascular network, the device captured VEGF-driven choroidal neovascularization through RPE barriers. Together, these establish the µSiM-OBRB as a platform for retinal drug bioavailability measurement, toxicity detection, and efficacy evaluation.

## Experimental

### µSiM device components and assembly

Nanoporous (NPSN, NPSN100.C-1LZ.0), microporous (MPSN, MPSN400-3L- 0.5HP, 0.5 µm diameter), and dual-scale (DSSN, DSSN100.C-1lZ.0-3.0A1, 3 µm micropore diameter) silicon nitride membrane chips were acquired from SiMPore Inc. (Henrietta, NY). NPSN membranes are approximately 100 nm thick with 35-65 nm pore diameters, total porosity of 9-27%, and membrane window area of 0.7 mm by 2 mm.

MPSN membranes have a thickness of 400 nm, a pore diameter of 0.5 µm, and a total porosity of 18-22%. MPSN membranes have three parallel membrane windows, each with dimensions of 0.7 mm by 2 mm. To create DSSN membranes, 3 µm micropores were etched onto a nanoporous membrane substrate, increasing the total porosity by 0.6-1%.^52^ The nanoporous background of DSSN membranes is approximately 100 nm thick, has pore diameters ranging from 35-80 nm, has a total porosity between 9-27%, and has a membrane window area of 0.7 mm by 2 mm. The Microfluidic device featuring a Silicon nitride Membrane (µSiM) chip comprises two main components. µSiM Component 1 (Aline, Inc.) is the upper portion of the device, which houses the NPSN membrane and an open-top well that contains a maximum 100 µL of media or buffer. µSiM Component 2 (Aline, Inc.), the bottom portion of the device, has a simple fluidic channel with a volume of 10 µL and a surface area of 44 mm^2^. µSiM Component 1 is made primarily of acrylic, and µSiM Component 2 is made primarily of PET with a cyclic olefin copolymer (COP) floor for imaging. Detailed dimensions and channel geometry for the µSiM device platform have been previously described.^46,53,54^

µSiM devices were assembled in sterile conditions in biosafety cabinets as previously described.^53^ Before assembly, µSiM Component 1 and Component 2 were rinsed with 70% ethanol and allowed to dry while being exposed to sterilizing UV for at least 15 minutes. NPSN membrane chips were bonded to the upper acrylic chip component using silicone-based pressure sensitive adhesive (PSA) and an aluminum alignment and assembly device. µSiM Component 2 was then bonded to µSiM Component 1 using a separate chip assembly device and PSA. Poly(dimethylsiloxane) (PDMS) flow bumper modules were added to fully assembled µSiM devices for fluidic experiments, including the fluidic small molecule permeability and retinal drug uptake assays to enable tight seals with fluidic circuits. To align the flow bumper module with the fluidic access ports of µSiM Component 1, blunt 21-gauge stainless steel dispenser cannulas (Jensen Global Inc., Santa Barbara, CA) were used. To adhere the flow bumpers to µSiM Component 1, PSA was applied and cured for at least 24 hours inside a 37 °C cell culture incubator.

### µSiM-OBRB ARPE-19 cell barrier cultures

ARPE-19 cells (CRL-2302, ATCC, Manassas, VA) between passages 5-9 were cultured using ARPE-19 growth media (ARPE-19-GM: Dulbecco’s Modified Eagle Medium: nutrient mixture F12 (DMEM/F12) + 10% fetal bovine serum (FBS) + 1% antibiotic-antimycotic). For µSiM-OBRB ARPE-19 cultures, fully assembled µSiM devices with NPSN membranes were pre-treated with 0.025 mg mL^−1^ laminin (ThermoFisher Scientific, Waltham, MA) for 24 hours prior to cell seeding. ARPE-19 cells were seeded in the apical well of µSiM chips at a density of 50,000 cells cm^−2^ and allowed to proliferate for the first 6 days of culture in growth medium, with media exchanged every other day. Starting on day 7, growth media was replaced with a modified ARPE-19 differentiation media (ARPE-19-DM: DMEM/F12 + 1% FBS + 10 mM Nicotinamide + 1% antibiotic-antimycotic).^55^ On days 7, 14, 21, or 28, µSiM-OBRB ARPE-19 monocultures were analyzed for transepithelial electrical resistance (TEER) and/or small molecule permeability then fixed using 4% paraformaldehyde (I28800, ThermoFisher, Waltham, MA).

### µSiM-OBRB ARPE-19 + HUVEC co-cultures

For ARPE-19 / human umbilical vein endothelial cell (HUVEC, Lonza, Basel, Switzerland) co-cultures, HUVECs were cultured and passaged using VascuLife® VEGF Endothelial Medium (Lifeline Cell Technologies, Frederick, MD) with 1% antibiotic-antimycotic. Before seeding HUVECs, the bottom channel of the µSiM device was coated with fibronectin at least 24 hours in advance to enable HUVEC adhesion to the bottom channel surface and the nanomembrane (0.175 mg mL^−1^ in DPBS). HUVECs were seeded in the bottom channel of µSiM chips at a volumetric cell density of 5,000,000 cells mL^−1^ of media 12 days after ARPE-19 cells were initially seeded. Immediately after HUVEC seeding, µSiM chips were inverted to allow HUVECs to adhere to the underside of NPSN membranes. Due to the small fluid volumes, surface tension prevented ARPE-19 cultures in the apical well from drying out. Vasculife was pipetted into the bottom channel to culture HUVECs, while ARPE-19-DM was added to the apical well to supplement the ARPE-19 cells, exchanging media daily until day 14, when ARPE-19 + HUVEC co-cultures were analyzed for small molecule permeability or the LIVE/DEAD assay then fixed using 4% paraformaldehyde.

### Assessment of transepithelial electrical resistance (TEER)

To measure TEER in µSiM-OBRB ARPE-19 + HUVEC co-cultures, a previously developed impedance module for µSiM devices and the associated method for measuring transepithelial electrical resistance (TEER) were used.^54,56^ Briefly, 21-gauge stainless steel cannulas were used as electrodes and attached to a Gamry Reference 600 potentiostat in a two-electrode configuration. One electrode was positioned in the apical well, and another was inserted into the bottom channel. Impedance spectra were acquired between a frequency range of 2.5 Hz to 1 MHz. TEER was reported in Ω cm^2^ as the impedance measured at 12.4 Hz.^54,56^ Before TEER measurements, the area of the NPSN membrane window (*A*) was measured using brightfield microscopy and ImageJ. The total systemic impedance was determined by measuring impedance of µSiM devices before cell culture (*R_0_*) and once again at the experimental endpoint after cell barrier formation (*R_t_*). *R_0_* was subtracted from *R_t_* then multiplied by the NPSN membrane window area (*A*) to determine systemic TEER (*TEER_Sys_*) using **Equation 1**. In parallel, µSiM devices without cells were maintained in identical conditions and measured at the same endpoints to determine *TEER_Control_*. To calculate the impedance of ARPE-19 tissue barriers (*TEER_Cells_*), the impedance of control devices *TEER_Control_* was subtracted from *TEER_Sys_* (**Equation 2**).

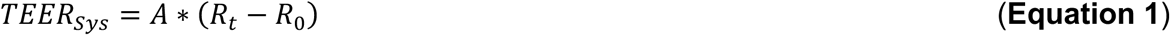

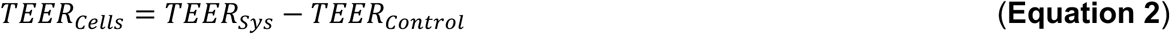

### Assessment of static small-molecule permeability

To measure small molecule permeability of µSiM devices, a previously described method was employed.^46^ 14 days (unless otherwise specified) after initial cell seeding, µSiM-OBRB devices were rinsed with ARPE-19 growth media in both channels prior to the permeability assay, and any bubbles were removed prior to starting the assay. 457 Da Lucifer Yellow (ThermoFisher, Waltham, MA) was mixed into ARPE-19 growth media at a concentration of 150 µg mL^−1^. The media in the apical well of µSiM devices was aspirated, then replaced with 100 µL of Lucifer Yellow media. The µSiM was then incubated at 37°C for 1 hour to allow the dye to diffuse. After the 1-hour incubation period, the lucifer yellow media was removed from the apical well, and a P200 pipette tip filled with 50 µL of ARPE-19-GM was added to one of the flow access ports of the µSiM device to serve as a temporary reservoir for the bottom channel. A second P200 pipette tip was used to extract a 50 µL media sample from the opposite flow port, which was then transferred to a black 96-well plate. To create a standard curve, a 2:1 dilution ladder made from the lucifer yellow media solution of known lucifer yellow concentrations was included in each plate. A microplate reader (Tecan Life Sciences, Mannendorf, Switzerland) was used to measure the fluorescence intensity (ex. 428 nm, em. 536 nm) in each well. Cell permeability (*P_Cells_*) was calculated using **Equations 3** and **4**, where *C_t_* is the concentration of lucifer yellow inside the bottom channel at time *t*, V is the media volume, *C_0_* is the initial concentration of lucifer yellow that was added to the apical well of the µSiM device, and *A* is the area of the membrane window. *P_Systemic_* is the measured systemic permeability of cell barriers and nanomembranes, and *P_Control_* is the permeability of a µSiM device without cells, which serves as a control.

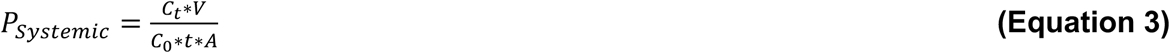

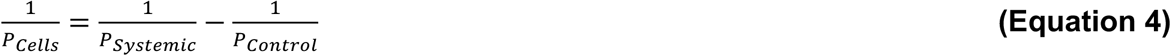

### Measurements of retinal drug transport under flow

Retinal uptake of several small molecule drug compounds was assessed in fluidic conditions. A peristaltic pump with a flow rate set to 100 µL min^−1^ was used to perfuse ARPE-19-GM mixed with lucifer yellow (150 µg mL^−1^), digoxin (200 ng mL^−1^), urea (960 µg mL^−1^), or the drug cocktail through the bottom channel of µSiM devices for 1 hour (**Fig. S1†**). To measure the concentration of lucifer yellow, each sample was added to a black 96-well plate, and a microplate reader (Tecan Life Sciences, Zurich, CH) was used to measure the fluorescence intensity (ex. 428 nm, em. 536 nm). To measure the concentration of digoxin in each media sample, a Human Digoxin ELISA Kit (Creative Diagnostics, Shirley, NY) was used. A colorimetric Urea Assay Kit III (Sigma-Aldrich, Burlington, MA) was used to measure urea concentration. The fluidic circuit was attached to the PDMS flow bumpers to minimize bubble formation and ensure uninterrupted flow. Ganciclovir, aztreonam, fluconazole, voriconazole, and rosiglitazone were combined into a drug cocktail (each at 1 µg mL^−1^ except voriconazole at 20 µg mL^−1^, in DMSO) and simultaneously perfused through the bottom channel of µSiM chips. Dynamic ranges for multiple small-molecule compounds were determined by preparing dilutions and measuring analyte concentrations by liquid chromatography-mass spectrometry (LC-MS). The apical well was sampled at *t* = 0 min, 30 min, and 60 min, with media replacement at each sampling step. After the retinal uptake assay, µSiM-OBRB ARPE-19 + HUVEC co-barrier cultures were fixed using 4% paraformaldehyde (PFA) and prepared for immunohistochemistry and confocal imaging. LC-MS was used to quantify the concentration and cumulative mass transport of each compound.

To prepare samples for LC-MS analysis, samples were diluted by a factor of 2, and 500 ng mL^−1^ of each of the following internal standards was added: fluconazole-d4, voriconazole-d3, aztreonam-d3, digoxin-d3, rosiglitazone-d3, and hydroxychloroquine- d4 (Cayman Chemical, Ann Arbor, MI). For hydrophobic interaction liquid chromatography (HILIC) LC-MS analyses, 90 µL of 90% LC-MS grade acetonitrile (A955, Fisher Scientific) was added to 10 µL media samples and incubated on ice for 30 minutes with vortexing. Following incubation, samples were centrifuged at 17,000 x g, 4 °C, for 10 minutes, and the supernatant was transferred to glass vials for analysis. For reverse phase LC-MS analyses, 30 µL of LC-MS grade acetonitrile was added to 10 µl media samples and incubated on ice for 30 minutes with vortexing. Following incubation, samples were centrifuged at 17,000 x g, 4 °C for 10 minutes, then 20ul of the supernatant was combined with 30 µL of LC-MS grade water. This solution was centrifuged at 17,000 x g, 4 °C for 10 minutes, and the supernatant was transferred to glass vials for analysis

For LC-MS analysis, samples were analyzed by high resolution mass spectrometry with an Orbitrap Exploris 240 (Thermo) coupled to a Vanquish Flex liquid chromatography system (Thermo). For both HILIC analysis and reverse phase analysis, mobile phase A was 100% LC-MS grade water with 10 mM ammonium formate and 0.125% formic acid. Mobile phase B was 90% acetonitrile with 10 mM ammonium formate and 0.125% formic acid. For HILIC analysis, 5 µL of samples were injected on a Waters XBridge XP BEH Amide column (150 mm length × 2.1 mm inner diameter, 2.5 µm particle size) maintained at 25C, with a Waters XBridge XP VanGuard BEH Amide (5 mm × 2.1 mm id, 2.5 µm particle size) guard column. The gradient was 0 minutes, 100% B; 2 minutes, 100% B; 3 minutes, 90% B; 5 minutes, 90% B; 6 minutes, 85% B; 7 minutes, 85% B; 8 minutes, 75% B; 9 minutes, 75% B; 10 minutes, 55% B; 12 minutes, 55% B; 13 minutes, 35%, 20 minutes, 35% B; 20.1 minutes, 35% B; 20.6 minutes, 100% B; 22.2 minutes, 100% B all at a flow rate of 150 µL min^−1^, followed by 22.7 minutes, 100% B; 27.9 minutes, 100% B at a flow rate of 300 µL min^−1^, and finally 28 minutes, 100% B at flow rate of 150 µL min^−1^, for a total length of 28 minutes. The H-ESI source was operated in positive mode at spray voltage 3500 with the following parameters: sheath gas 35 au, aux gas 7 au, sweep gas 0 au, ion transfer tube temperature 320 C, vaporizer temperature 275°C, MS1 mass resolution of 120,000 FWHM, RF lens at 70%, standard automatic gain control (AGC), MS2 fragmentation with normalized collision energies of 20%, 30%, 50%, 75%, 100% and resolution at 15,000 FWHM. For reverse phase analysis, 5 µL of samples were also injected on a Thermo Accucore column (100 mm length × 2.1 mm id, 2.6 µm particle size) maintained at 40C. The gradient was 0 minutes, 15% B; 1 minutes, 15% B; 3 minutes, 25% B; 9 minutes, 65% B; 13 minutes, 100% B; 15 minutes, 100% B; 15.1 minutes, 15% B; and finally, 20 minutes, 15% B at flow rate of 400 µL min^−1^, for a total length of 20 minutes.

The H-ESI source was operated in positive mode at spray voltage 3500 with the following parameters: sheath gas 50 au, aux gas 10 au, sweep gas 1 au, ion transfer tube temperature 325 C, vaporizer temperature 300 C, MS1 mass resolution of 120,000 FWHM, data dependent MS2 (ddMS2) fragmentation with normalized collision energies of 20%, 30%, 40%, and MS2 resolution at 15,000 FWHM. LC-MS data were analyzed by Maven software for peak determination and annotation via matching to external standards. Peak values of Voriconazole, Rosiglitazone, Fluconazole, and Aztreonam in experimental samples and calibration samples were normalized to those of Voriconazole-d3 (3 ng mL^−1^), Rosiglitazone-d3 (0.5 ng mL^−1^), Fluconazole-d4 (3 ng mL^−1^), and Aztreonam-d6 (6 ng mL^−1^) respectively, and calibration curves were plotted using linear regression for quantitation.

**Equation 5** was used to calculate the retinal uptake index (RUI) for a specific molecule, such that Q_t_ is the cumulative concentration of a given compound that in the retinal chamber over time and Q_0_ is the initial compound concentration in the endothelial channel media. To calculate the RUI for cell barriers (*RUI_Cells_*), the baseline RUI of devices without cells (*RUI_Control_*) was subtracted from the Systemic RUI of µSiM chips with ARPE-19 cells and HUVECs (*RUI_Systemic_*) (**Equation 6**). SDF files downloaded from the PubChem database were inputted into ADMETLab 3.0 to predict Log P and Log D_7.4_ values for each small molecule drug compound.^57–59^

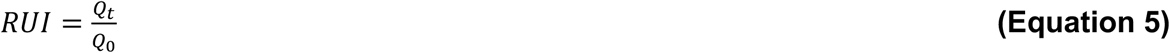

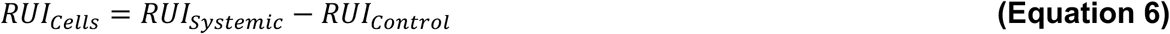

### Digoxin toxicity assay

Digoxin was introduced into ARPE-19-GM at a concentration associated with clinical digoxin toxicity (10 ng mL^−1^).^60–62^ Digoxin media was added to µSiM-OBRB ARPE-19 + HUVEC co-barrier cultures starting on day 12 and maintained for 48 hours. On day 14, the small-molecule permeability assay was performed using lucifer yellow as described above. After the permeability assay, µSiM-OBRB ARPE-19 + HUVEC co- barrier cultures were incubated with a LIVE/DEAD Cell Imaging Kit (488/570) (Invitrogen, Waltham, MA) for 15 minutes, then imaged using confocal microscopy. Live cells were stained with Calcein AM, and dead cells were stained with BOBO-3 Iodide. ImageJ was used to quantify cell viability. After imaging with the LIVE/DEAD assay, µSiM-OBRB ARPE-19 + HUVEC co-barrier cultures were fixed using 4% PFA.

### Forming 3D microvasculature in µSiM-OBRB using injectable fibrin hydrogels

Human umbilical vein endothelial cells (HUVECs, Lonza, Basel, Switzerland) between passages 2-5 were used due to their ubiquitous usage as a model endothelial cell type. HUVECs were cultured and passaged using VascuLife® VEGF Endothelial Medium (LL-0003, Lifeline Cell Technologies, Frederick, MD) supplemented with 1% antibiotic-antimycotic (15240062, ThermoFisher). Human mesenchymal stem cells (hMSCs) were derived from bone marrow aspirates as previously described.^63,64^ hMSCs were cultured and passaged using low glucose DMEM (Dulbecco’s Modified Eagle Medium, low glucose, pyruvate, 11885084, ThermoFisher) supplemented with 10% FBS (A5670701, ThermoFisher) and 1% antibiotic-antimycotic (15240062, ThermoFisher).

To prepare injectable fibrin hydrogels, a 10 mg mL^−1^ solution of fibrinogen (341578-500MG, Millipore Sigma, Burlington, MA) was dissolved in Dulbecco’s phosphate buffered saline (DPBS, 14190144, ThermoFisher Scientific) and kept at 37 °C overnight. The following day, the fibrinogen solution was centrifuged at 200 rcf for 2 minutes, then filtered using a 0.22 µm low protein-binding syringe filter (SLGPM33RS, Millipore Sigma, Burlington, MA). A thrombin solution comprised of 19.2 µL of DPBS and 0.8 µL of 10 U mL^−1^ thrombin (605195, Millipore Sigma, Burlington, MA) was prepared.

After passaging, HUVECs and hMSCs were mixed in the thrombin solution at a 1:1 ratio, each with a density of 2.5 × 10^6^ cells mL^−1^ for each cell type, resulting in a final cell concentration of 5 × 10^6^ cells mL^−1^. Before mixing the fibrin solution, any media in the apical well was aspirated from µSiM-OBRB devices to prevent gel dilution, and bubbles were removed from the bottom channel. For each device, the fibrinogen solution was mixed with the cell suspension/thrombin solution and rapidly mixed to create a 40 µL precursor solution, with final fibrinogen concentration of 5 mg mL^−1^ and 2 U mL^−1^ thrombin.^65,66^ The 10 µL of the precursor solution was immediately injected into the bottom channel of a µSiM-OBRB device using a P20 micropipette, then allowed to polymerize for 30 minutes at 37 °C and 5% CO_2_ before Vasculife was added to the apical well. Vasculife was exchanged in the apical well of µSiM-OBRB HUVEC + hMSC co-cultures every alternating day for up to 14 days. Additional cell seeding ratios were tested in 96-well plates, including 1:1 HUVECs:hMSCs, 2:1 HUVECs:hMSCs and 3:1 HUVECs:hMSCs, all at a final cell concentration of 5 × 10^6^ cells mL^−1^.

### Modeling choroidal neovascularization using µSiM-OBRB ARPE-19 + HUVEC + hMSC tri-cultures

To model choroidal neovascularization in µSiM-OBRB, DSSN membranes, which feature large micropores that enable endothelial cell transmigration, were used.^52,67,68^ µSiM-OBRB devices with MPSN membranes that prevent cellular transmigration were included as a negative control. Starting on day 10, µSiM-OBRB tri-cultures were treated with ARPE-19-DM with 25 ng mL^−1^ vascular endothelial growth factor A (VEGF-A) supplementation to induce apical endothelial cell migration. VEGF-supplemented ARPE-19-DM was exchanged daily until day 14, when VEGF+ µSiM-OBRB ARPE-19 + HUVEC + hMSC tri-cultures were fixed using 4% paraformaldehyde before antibody staining. To track endothelial cell vascular invasion, Z-stack confocal images were acquired, and CD31+ HUVECs that migrated above ARPE-19 cells were identified in orthogonal projections of confocal images.

### Quantification of vessel parameters based on confocal microscopy images

For vasculogenesis-emulating conditions, vessel positive areas of images were identified using endothelial cell markers CD31 or VE-Cadherin. To quantify vascular parameters for Z-stack confocal images of microvascular networks in vascularized µSiM-OBRB devices, average vessel diameter was measured by using Fiji with the Vessel Analysis plug-in package (diameter), and AngioTool was used to measure density, network length, branching points, and average branch length.^69,70^ For angiogenesis analyses, Z-stack images were quantified using the ImageJ “Sprout Morphology” plug-in to quantify cell sprouting in hydrogels. Cell trackers were used to distinguish HUVECs and hMSCs, respectively.

### Immunohistochemistry, confocal microscopy, and image quantification

µSiM-OBRB ARPE-19 monocultures, ARPE-19 + HUVEC co-cultures, and µSiM- OBRB ARPE-19 + HUVEC + hMSC tri-cultures were fixed using 4% paraformaldehyde. A blocking and staining buffer containing 0.25% Triton X-100 / 1% bovine serum albumin in Dulbecco’s Phosphate Buffer Saline (DPBS) was used to permeabilize cells and prevent non-specific binding. The apical chamber and basal channel of each µSiM chip was incubated with primary, secondary, and pre-conjugated antibodies in separate steps overnight at 4°C. Between each staining step, µSiM devices were each rinsed at least 3 times with the staining buffer and covered from light with aluminum foil. After all staining steps were completed, the apical well and bottom channel of each µSiM device was rinsed at least 3 times with DPBS.

Primary antibodies that were used include: Rabbit IgG RPE65 Polyclonal antibody (ThermoFisher Scientific), sheep IgG anti-CD31 (R & D Systems), mouse IgG2a anti-α-SMA (Invitrogen), mouse IgG anti-VE-Cadherin (Santa Cruz Biotechnology), rabbit IgG anti-Ezrin (Cell Signaling Technology). Secondary antibodies used include goat IgG anti-mouse AlexaFluor 488 (Invitrogen), goat IgG anti-mouse AlexaFluor 568 (Invitrogen), goat IgG anti-Rabbit AlexaFluor 488 (Invitrogen), goat IgG anti-rabbit AlexaFluor 568, and goat IgG anti-Rabbit AlexaFluor 647 (Invitrogen). Pre-conjugated antibodies used include mouse IgG anti-Collagen IV AlexaFluor 647 (Invitrogen), mouse IgG anti-CD31 monoclonal antibody (MEM-05) AlexaFluor 488 (ThermoFisher Scientific), mouse IgG recombinant mouse monoclonal antibody (ZO1- 1A12), AlexaFluor Plus 555 (ThermoFisher Scientific). A full list of antibodies used for this study is tabulated in **Table S2†**.

A Dragonfly spinning disk confocal microscope was used to acquire Z-stack fluorescence images of each µSiM device at 10X and 40X magnifications. The Z-step sizes were 1.33 µm for images acquired at 10X magnification and 0.5 µm for images acquired at 40X magnification. Up to four channels were acquired per image. Maximum projections of Z-stack confocal images were analyzed using Fiji (ImageJ) or Imaris. Orthogonal projections of Z-stack images were analyzed using Imaris.

### Statistical analyses

All data were analyzed using Graphpad Prism and Microsoft Excel. Student’s t-test was used to compare means in experiments with two groups. One-way ANOVA was used to analyze differences in variance, and Tukey’s honest significance test for multiple comparisons was used to compare the means in experiments with three or more groups. Data plotted in figures are presented as mean ± standard deviation (SD), with N indicating the number of independent devices. Where a single value is reported in the text or in a table, it is given as mean ± standard error of the mean (SEM) and labeled (SEM), so that the error accompanying a reported benchmark value is the error in that value.

## Results

### µSiM-OBRB ARPE-19 monocultures form functionally tight barriers with a mature phenotype

ARPE-19 cells were seeded in the apical well of µSiM devices on top of laminin-coated NPSN membranes and matured using Nicotinamide-supplemented differentiation media (**Fig. 2A, B**, **Fig. S2†**).^55^ Immunohistochemistry was used to study tissue development in µSiM-OBRB devices. Zonula occludens-1 (ZO-1) was stained to assess cellular tight junction expression and hexagonal cobblestone morphology.^71,72^ µSiM-OBRB ARPE-19 cells formed confluent monolayers with ZO-1+ tight junctions and hexagonal packing (**Fig. 2C, D**). Ezrin, a structural protein that localizes to apical microvilli in RPE, was used as a marker for cell polarization.^72,73^ Collagen IV was also stained as a marker of the RPE basement membrane.^74–76^ Orthogonal projections of Z-stack confocal images of µSiM-OBRB ARPE-19 cultures show that Ezrin is apically localized relative to cell nuclei and ZO-1 (**Fig. 2E**). Furthermore, Collagen IV was basally deposited by µSiM-OBRB ARPE-19 cells, mimicking the maturation of basal lamina (**Fig. 2F**). These data show that µSiM-OBRB ARPE-19 form polarized barrier tissues with tight junction expression and morphologically mature hexagonal packing.

**Fig. 2.**
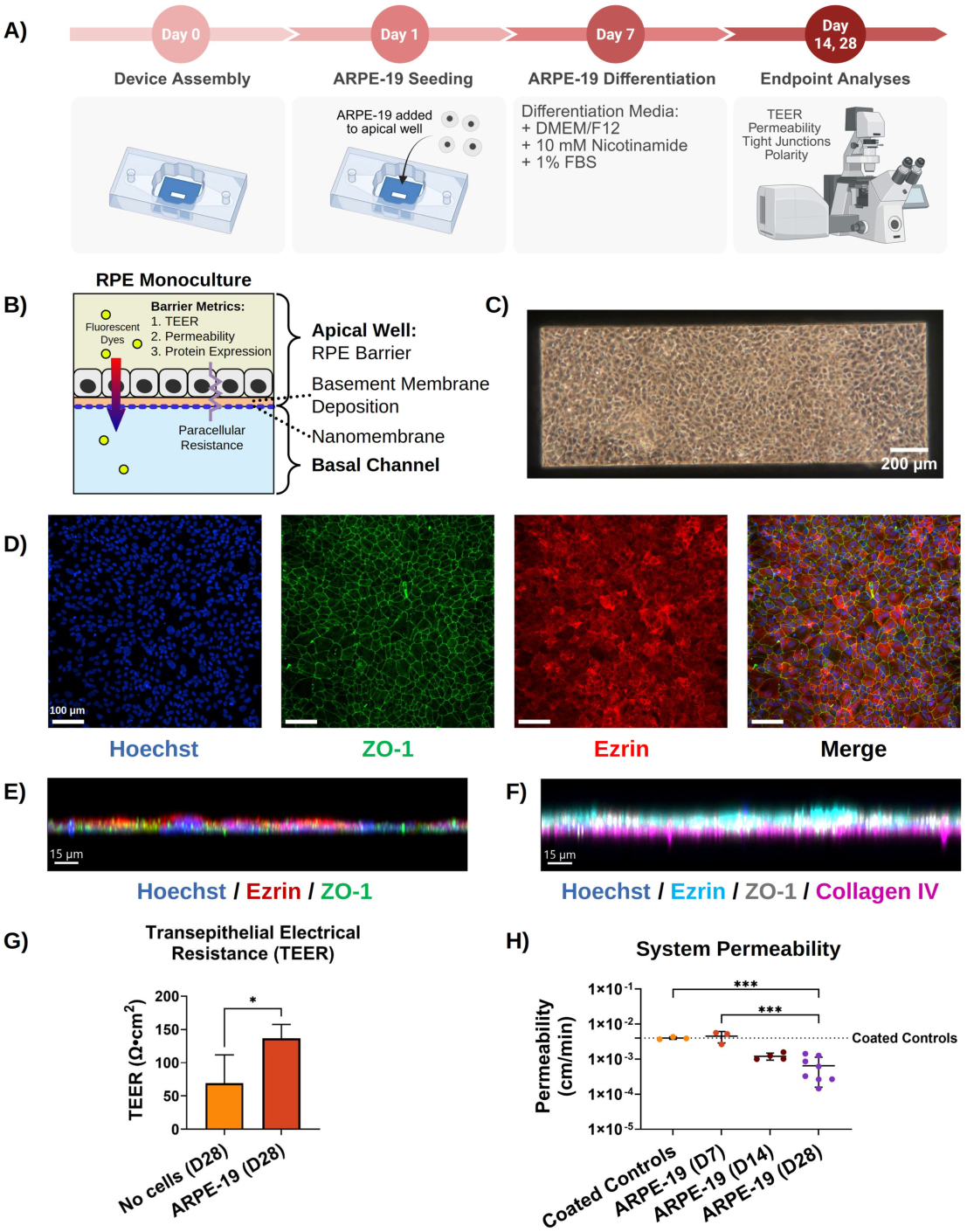
µSiM-OBRB ARPE-19 monocultures form tight barriers, as measured by transepithelial electrical resistance (TEER) and small molecule permeability. **(A)** Protocol for seeding ARPE-19 monoculture barriers on µSiM-OBRB devices. Made with Biorender. **(B)** ARPE-19 were seeded on the top surface of NPSN membranes in µSiM- OBRB devices. To assess barrier function, TEER and small molecule permeability were measured. **(C)** Phase microscopy image of ARPE-19 cells on a NPSN membrane. Image taken on day 14. **(D)** Confocal Z-stack max projection images of ARPE-19 cells seeded on top of a NPSN membrane. Blue: Hoechst, Green: ZO-1, Red: Ezrin. Device fixed on day 21. **(E)** Ezrin (red) is apically localized compared to ZO-1 (green) and cell nuclei (blue), indicating cell polarization. Device fixed on day 28. **(F)** µSiM-OBRB ARPE- 19 cells deposit basal collagen IV (magenta) below apical ezrin (cyan) and ZO-1 (white), mimicking Bruch’s membrane deposition. Device fixed on day 21. **(G)** TEER after 28 days. ARPE-19 monolayers roughly double the resistance of the cell-free coated device (error bars represent SD; values reported in the Results), confirming a functional barrier. N = 5 (ARPE-19), N = 3 (no cells). Statistics: unpaired t-test, *p < 0.05. **(H)** Small molecule permeability assays confirm that both ARPE-19 monocultures form tight barriers that restrict the apicobasal diffusion of small molecules under static conditions (Lucifer Yellow, 457 Da). Mean ± SD. N=3-8. Statistics: one-way ANOVA with Tukey’s Multiple Comparison’s test. *p<0.05, **p<0.005, ***p<0.0005.

Transepithelial electrical resistance (TEER) reports the ionic conductance of the barrier, which is dominated by the paracellular pathway through tight junctions.^54,56,77^ After 28 days, TEER of µSiM-OBRB ARPE-19 barriers was measured and compared to µSiM devices maintained in culture media but without cells.^54,56^ Subtracting day 0 values from both, the TEER of ARPE-19 monocultures in µSiM-OBRB increased by 137 ± 9 Ω cm^2^, however control chips also increased over this time: 69 ± 25 Ω cm^2^ (**Fig. 2G, Fig. S3†**). Subtracting these values provides an estimated cellular TEER of 68 ± 26 Ω cm^2^ (mean ± SEM, n = 5 devices vs. n = 3 cell-free controls; unpaired t-test, p = 0.02).

To assess the small molecule permeability of µSiM-OBRB ARPE-19 cultures, lucifer yellow was incubated for 1 hour in the apical well of µSiM-OBRB ARPE-19 cultures.^46^ Laminin coated µSiM device controls without cells were used to establish a baseline for both TEER and permeability measurements. Small molecule permeability for µSiM-OBRB ARPE-19 cultures was assessed on days 7, 14, and 28. The permeability of 7-day µSiM-OBRB ARPE-19 cultures was similar to coated controls. Compared to controls, the permeability of 14-day µSiM-OBRB ARPE-19 cultures was about 3-fold lower. 28-day µSiM-OBRB ARPE-19 cultures had the lowest permeability, over 6-fold lower than controls, and about 2-fold lower than 14-day cultures. (Controls: 3.98 ± 0.2 × 10^−3^ cm min^−1^, 7-day ARPE-19: 4.51 ± 0.9 × 10^−3^ cm min^−1^, 14-day ARPE- 19: 1.21 ± 0.1 × 10^−3^ cm min^−1^, 28-day ARPE-19: 6.48 ± 1.7 × 10^−4^ cm min^−1^) (**Fig. 2H**). Altogether, these data suggest that µSiM-OBRB supports the formation of tight and functional RPE barrier tissues (**Table 1**).

**Table 1.** Reported / estimated ARPE-19 barrier parameters in OBRB-Chip and transwell based in vitro models. A complete overview of reported TEER values for ARPE-19 cells cultured on Transwells is available in the supplemental (Table S3†)

| **ARPE-19 Barrier Property** | **Expected Value** | **$\mu\text{SiM-OBRB}$** |
| --- | --- | --- |
| Hexagonal Morphology | ZO-1 Tight Junctions<br>Hexagonal Packing | See Fig. 2C, D |
| Ezrin Polarization | Apical Ezrin Localization | See Fig. 2E |
| Bruch's Membrane Protein Deposition | Collagen IV, VI, laminin, fibronectin, elastin expression | See Fig. 2F |
| TEER ($\Omega \text{ cm}^2$) | 40–50 $\Omega \text{ cm}^2$ <sup>78–84</sup> (**Table S3†**) | $68 \pm 26 \Omega \text{ cm}^2$ (Fig. 2G) |
| Permeability ($\text{cm min}^{-1}$) | $7.8 \pm 2.5 \times 10^{-5}$ <sup>85</sup> to $1.6 \times 10^{-3} \text{ cm min}^{-1}$ <sup>86</sup> | $6.48 \pm 1.7 \times 10^{-4} \text{ cm min}^{-1}$ (Fig. 2H) |

### ARPE-19-HUVEC co-cultures form tight barriers and restrict apicobasal small-molecule transport

To further recapitulate the interface between the retinal pigment epithelium and the choriocapillaris, ARPE-19 cells and human umbilical vein endothelial cells (HUVECs) were seeded on opposite sides of nanomembranes in µSiM-OBRB devices (**Fig. 3A–D**). To form the ARPE-19-HUVEC interface, HUVECs were seeded in the bottom channel 12 days after ARPE-19 cells were seeded, and µSiM-OBRB devices were inverted overnight for HUVEC adherence to the fibronectin-coated underside of the nanomembrane similar to methods described for µSiM-blood brain barrier models.^53^ ARPE-19 cells were stained with RPE65 or Ezrin, specific markers for RPE cells, and HUVECs were stained for CD31, an endothelial cell marker, to confirm direct juxtaposition (**Fig. 3C, D**). The 100 nm thinness of the nanomembrane makes it optically invisible in orthogonal projections of confocal Z stack images. HUVECs formed a monolayer that covered the inner surface of the basal fluidic channel. ZO-1 staining was used to assess tight junction formation by both ARPE-19 and HUVECs (**Fig. 3C**).

**Fig. 3.**
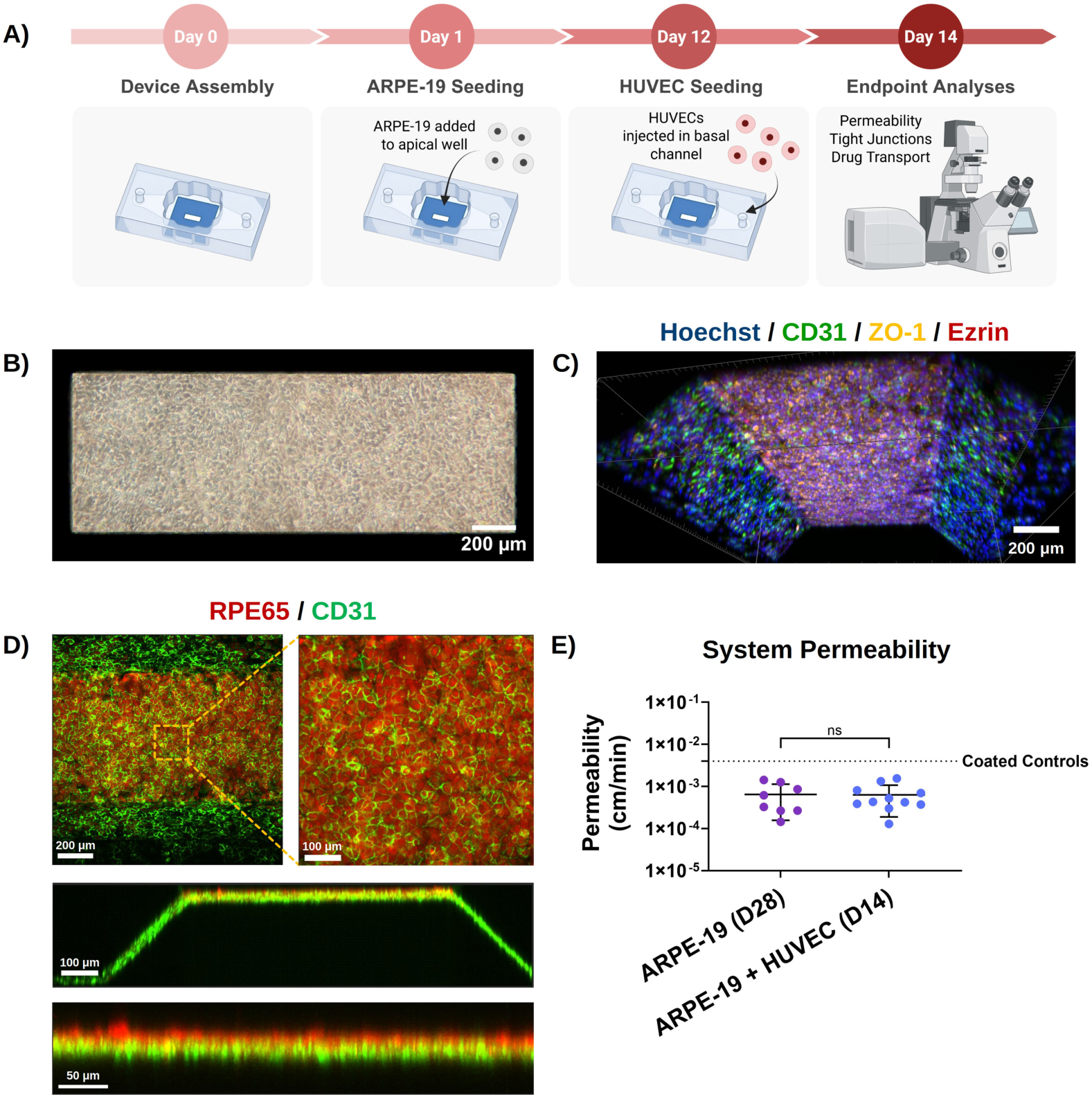
*ARPE-19 + HUVEC co-cultures form tight co-barriers on µSiM-OBRB chips.* **(A)** Protocol for co-culturing ARPE-19 + HUVEC co-barriers on µSiM-OBRB chips. Made with Biorender. **(B)** Phase microscopy image of ARPE-19 cells and HUVECs co-cultured on a NPSN membrane (Day 14). Scale bar: 200 µm. **(C)** 3D view of a Z-stack confocal image of µSiM-OBRB ARPE-19 + HUVEC co-cultures. Blue: Hoechst, Green: CD31, Yellow: ZO-1, Red: Ezrin. Scale bar: 200 µm. Device fixed on day 14. **(D)** Z-stack confocal images and orthogonal projections depicting µSiM-OBRB ARPE-19 + HUVEC co-cultures. ARPE-19 cells were stained RPE65 (red), and HUVECs were stained for CD31 (green). Orthogonal projections show that ARPE-19 are adhered to the top surface of the NPSN membrane, while HUVECs form a confluent monolayer on the surface of the lower channel under the nanomembrane. Scale bars: 200 µm, 100 µm, 50 µm. Image acquired on day 14. **(E)** Systemic permeability data for 28-day ARPE-19 monocultures and 14-day ARPE-19 + HUVEC co-cultures. The addition of HUVECs to 14-day ARPE-19 cultures lowers permeability and makes it comparable to 28-day ARPE-19 monocultures (Dye: Lucifer Yellow, 457 Da). Mean ± SD. N=8-11. Statistics: Student’s t-test *p<0.05, **p<0.005, ***p<0.0005, ****p<0.00005.

To assess the permeability of ARPE-19 + HUVEC co-barriers, lucifer yellow was used to measure the small molecule permeability of µSiM-OBRB ARPE-19 + HUVEC co-cultures after 14 days of culture (**Fig. 3E**). Despite the shorter culture duration, the addition of HUVEC barriers in the bottom channel improved barrier function; 14-day ARPE-19 + HUVEC co-cultures formed tight barriers with similar permeability to 28-day ARPE-19 monocultures (28-day ARPE-19 monocultures: 6.48 ± 1.7 × 10^−4^ cm min^−1^, 14-day ARPE-19 + HUVEC co-cultures: 6.34 ± 1.3 × 10^−4^ cm min^−1^) (mean ± SEM). On this basis, 14 days of co-culture was adopted as the standard barrier maturation period for all subsequent co-culture experiments, improving experimental throughput by halving the time to a usable barrier compared to monoculture alone.

### µSiM-OBRB ARPE-19 + HUVEC co-barriers restrict the retinal uptake of small-molecule drug compounds during flow

To model the retinal transport of small-molecule drugs under physiological conditions, a microfluidic circuit was connected to the bottom channel of µSiM-OBRB ARPE-19 + HUVEC co-cultures. A peristaltic pump set to a volumetric flow rate of 100 µL min^−1^ was used to perfuse the chip (**Fig. S1, S4†**). Based on COMSOL simulations, the wall shear stress at the membrane interface ranged from approximately 0.15 to 0.25 dyn cm^−2^ (**Fig. S5†**). In a preliminary evaluation of retinal uptake of small molecules, lucifer yellow, digoxin, and urea were added to ARPE-19 growth medium and perfused through the bottom channel for 1 hour with transport to the apical well measured by different means (see methods) at 0, 30, and 60 minutes. The retinal uptake index (RUI) for each small molecule was calculated as the ratio of diffused drug to the total supplied amount (**Equation 5**). ARPE-19 + HUVEC co-barriers reduced the cumulative transport of all three compounds (**Fig. S6†**).

To more rigorously evaluate the ability of the µSiM-OBRB to predict bioavailability, we tested a panel of five FDA-approved small molecules spanning more than four orders of magnitude in lipophilicity. Previous work has shown that the outer blood-retinal barrier is more permeable to compounds with higher lipophilicity than to hydrophilic compounds *in vivo*.^78,79^ To determine if µSiM-OBRB replicates this physiologic sorting, the molecular transport of drugs with a range of lipophilicities was measured. Specifically, a cocktail containing aztreonam, ganciclovir, fluconazole, rosiglitazone, and voriconazole was mixed into ARPE-19 growth media, then perfused through the basal channel of µSiM-OBRB chips for 1 hour (**Fig. 4A**). To measure the concentration of accumulated drug compounds in the apical well, liquid chromatography mass spectrometry (LC-MS) was used at 0 min, 30 min, and 60 min (**Fig. 4B**). Log P, the octanol-water partition coefficient for the non-ionized form of a given compound, and Log D_7.4_, the distribution coefficient of a given compound between n-octanol and a pH 7.4 buffer (most relevant to biological systems) were both used to represent lipophilicity (**Table 2**).^80^ Linear correlations between RUI and both Log D_7.4_ and Log P were observed with lipophilic compounds crossing ARPE-19 + HUVEC co-barriers more easily than hydrophilic compounds (**Fig. 4B–D, Fig. S7†**). For example, the RUI for ganciclovir was about 9-fold lower compared to rosiglitazone (ganciclovir: -0.19 ± 0.15, rosiglitazone: -0.02 ± 0.04).

**Fig. 4.**
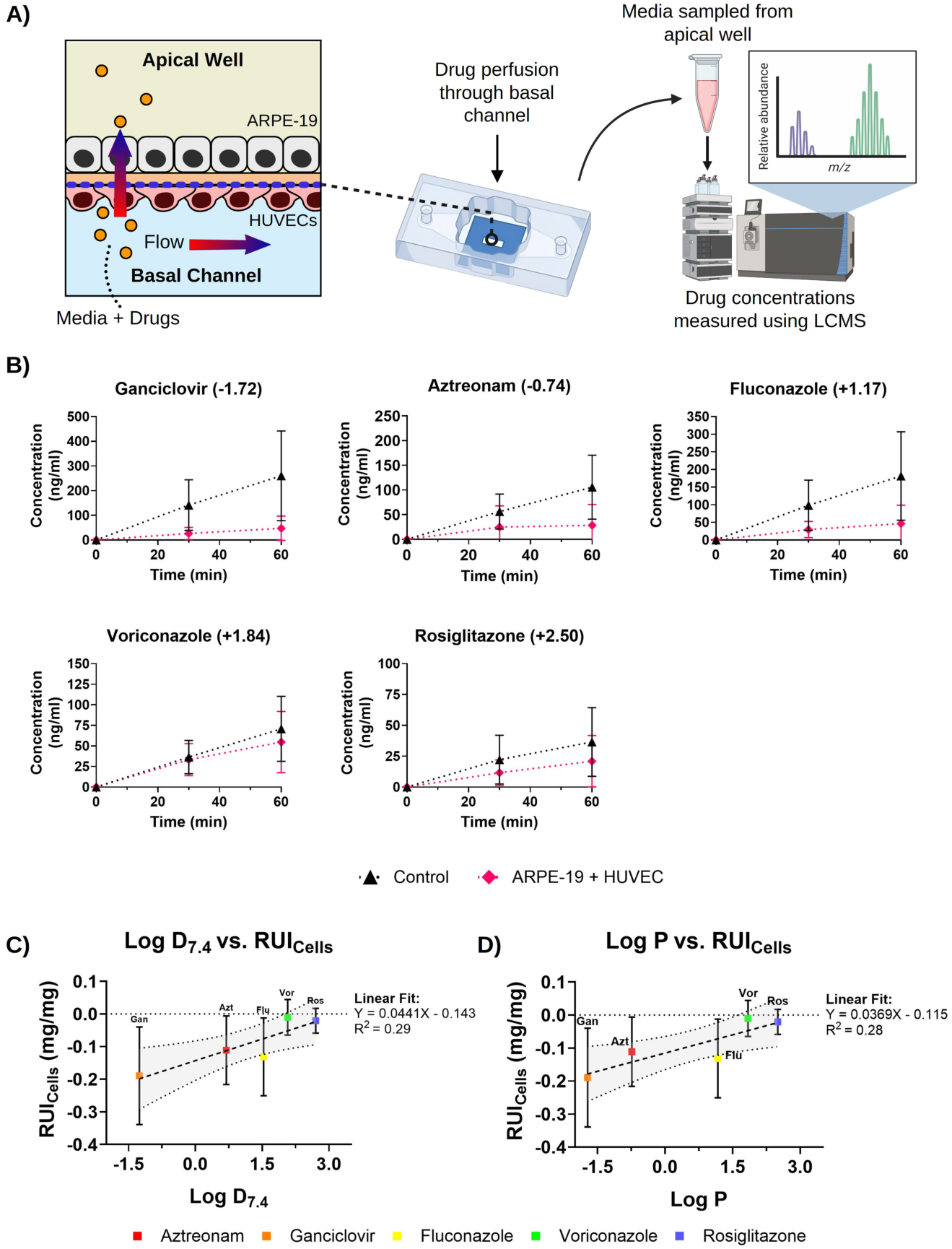
*ARPE-19 + HUVEC co-barriers reduce the retinal transport of small molecules.* **(A)** Small molecule drug compounds were perfused through the bottom channel of µSiM-OBRB for 1 hour, and media was sampled from the top well at 0, 30, and 60 min. Drug concentrations in media samples were quantified using liquid chromatography mass spectrometry. **(B)** Cumulative uptake of small molecule drug compounds in µSiM- OBRB with (magenta) or without (black) ARPE-19 + HUVECs barriers. Mean ± SD. N=4-5. **(C)** Cellular retinal uptake index (RUI) plotted against Log D_7.4_ for each drug compound. Dashed line: linear regression for LC-MS data, with Y = 0.0441X – 0.143, R^2^=0.29. Dotted lines and shaded area: 95% confidence interval range for the linear regression. N=4-5. **(D)** Cellular Retinal uptake index (RUI) plotted against Log P for each drug compound. Dashed line: linear regression for LC-MS data, with Y = 0.0369X – 0.115, R^2^=0.28. Dotted lines and shaded areas: 95% confidence interval range for the linear regression. Mean ± SD. N=4-5.

**Table 2.** The five FDA-approved small-molecule drugs of the LC-MS panel, with molecular weight, lipophilicity (represented by Log D7.4 or Log P), transport mechanism, assessment method, and references. ADMETLab 3.0 and the PubChem database were used to estimate Log P and Log D7.4 values 57,59,87.

|  |  |  |  |  |  |  |
| --- | --- | --- | --- | --- | --- | --- |
| **Compound** | **Molecular weight (Da)** | **Log P** | **Log $D_{7.4}$** | **Measurement Method** | **Transport Type** | **References** |
| Ganciclovir | 255.20 | -1.72 | -1.26 | LC-MS | Passive | <sup>88</sup> |
| Aztreonam | 435.44 | -0.74 | +0.69 | LC-MS | Passive | <sup>89</sup> |
| Fluconazole | 306.27 | +1.17 | +1.53 | LC-MS | Passive | <sup>90</sup> |
| Voriconazole | 349.31 | +1.84 | +2.07 | LC-MS | Passive | <sup>91</sup> |
| Rosiglitazone | 357.43 | +2.50 | +2.70 | LC-MS | Unknown | <sup>92–94</sup> |

### Digoxin causes loss of barrier integrity due to cell death in µSiM-OBRB ARPE-19 + HUVEC co-cultures

To model digoxin-induced retinal toxicity using µSiM-OBRB, ARPE-19 + HUVEC co- barrier cultures were treated with 10 ng mL^−1^ digoxin for 48 hours after 14 days of barrier development.^60,62,95^ Digoxin treatment induced major cell loss and caused the overall cell coverage of the membrane to decrease drastically, resulting in clear gaps in the cell barrier (**Fig. 5A**). The Calcein AM+ live cell area of digoxin-treated ARPE-19 + HUVEC co-cultures was reduced by about 50% compared to untreated ARPE-19 + HUVEC co-cultures (digoxin-treatment: 46 ± 14%, untreated: 90 ± 8%) (**Fig. 5A, B**). The ratio of live cell area / dead cell area was about 50-fold higher for untreated µSiM-OBRB cultures compared to digoxin treated chips (digoxin-treated: 13 ± 8, untreated: 650 ± 3, geometric mean ± geometric SD) (**Fig. 5B**). Digoxin-treated µSiM-OBRB cultures had about 7-fold as many dead cells compared to untreated controls (digoxin-treated: 176 ± 2, untreated: 25 ± 3, geometric mean ± geometric SD) (**Fig. S8†**). While digoxin-treated ARPE-19 + HUVEC co-barriers were still less permeable than control chips without cells, they were over 2-fold more permeable than untreated co-barriers (**Fig. 5C**), indicating that barrier integrity was significantly weakened (digoxin-treated: 1.41 ± 0.2 × 10^−3^ cm min^−1^, untreated: 6.34 ± 1.3 × 10^−4^ cm min^−1^).

**Fig. 5.**
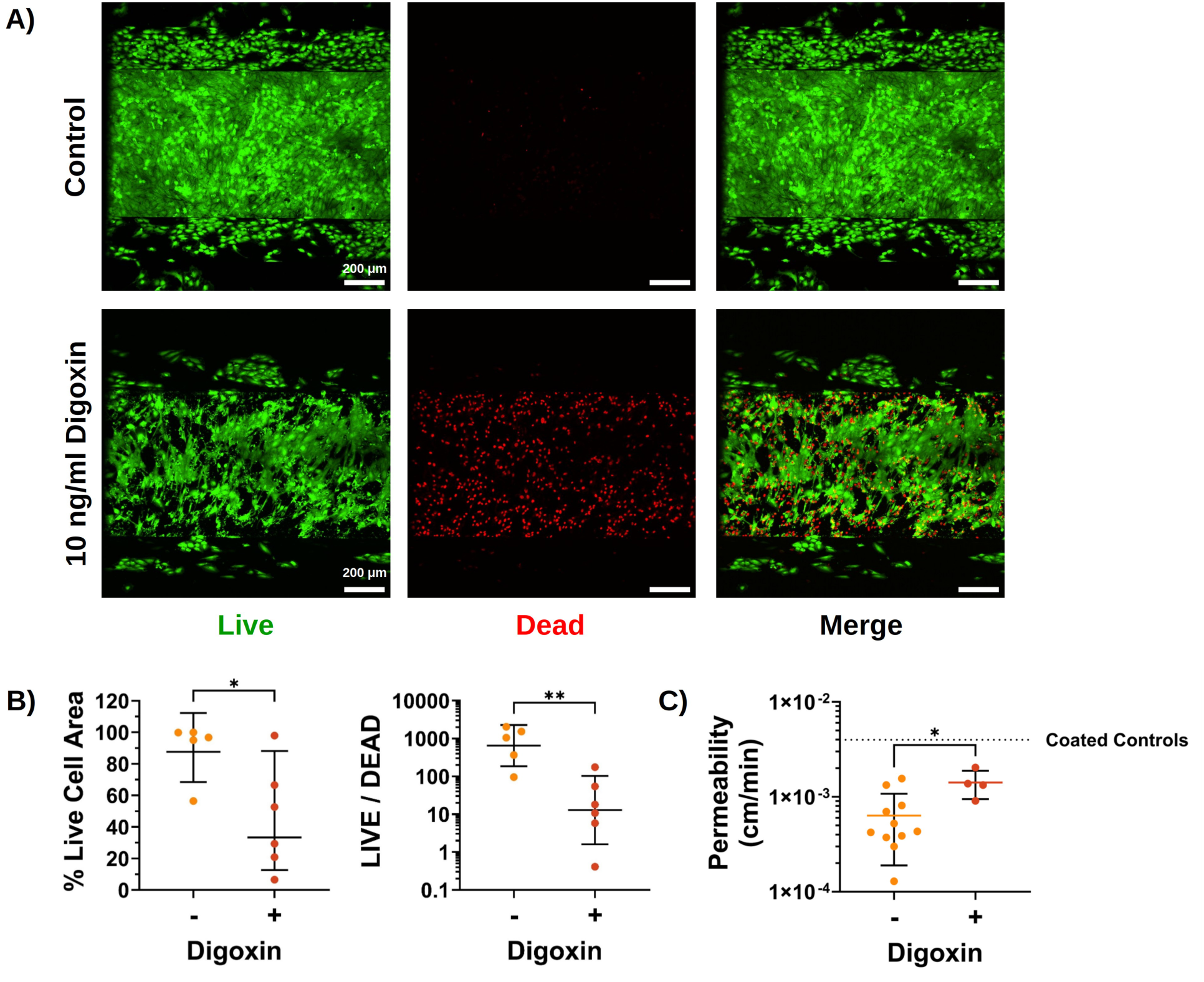
*Digoxin causes cell death and loss of barrier function in µSiM-OBRB ARPE-19 + HUVEC co-barrier cultures.* **(A)** Confocal Z-stack max projection images of µSiM-OBRB ARPE-19 + HUVEC co-cultures stained with the LIVE/DEAD cell imaging kit. Digoxin treatment caused cell death, loss of tight junctions and monolayer coverage. Images acquired on day 14. Green: live cells, Red: dead cells. **(B)** Quantification of LIVE/DEAD assay, showing that digoxin treatment caused significant cell death compared to untreated controls. Geometric mean ± SD. N=5-6. **(C)** Digoxin treatment caused µSiM- OBRB ARPE-19 + HUVEC co-cultures to be significantly more permeable to small molecule diffusion compared to controls, indicating loss of barrier function. Mean ± SD. N=4-11. Statistics: one-way ANOVA with Tukey’s Multiple Comparison’s test. *p<0.05, **p<0.005, ***p<0.0005, ****p<0.00005.

### Forming microvascular networks in µSiM-OBRB that mimic the choriocapillaris

HUVECs and hMSCs co-encapsulated in injectable fibrin hydrogels formed CD31+ and CD144+ microvascular networks in the bottom channel of µSiM-OBRB (**Fig. 6A, B, Fig. 7A–C**). Fibrin hydrogels with equal numbers of HUVECs and hMSCs supported the development of microvascular networks that most closely resembled the native choriocapillaris (**Fig. S9†**).

**Fig. 6.**
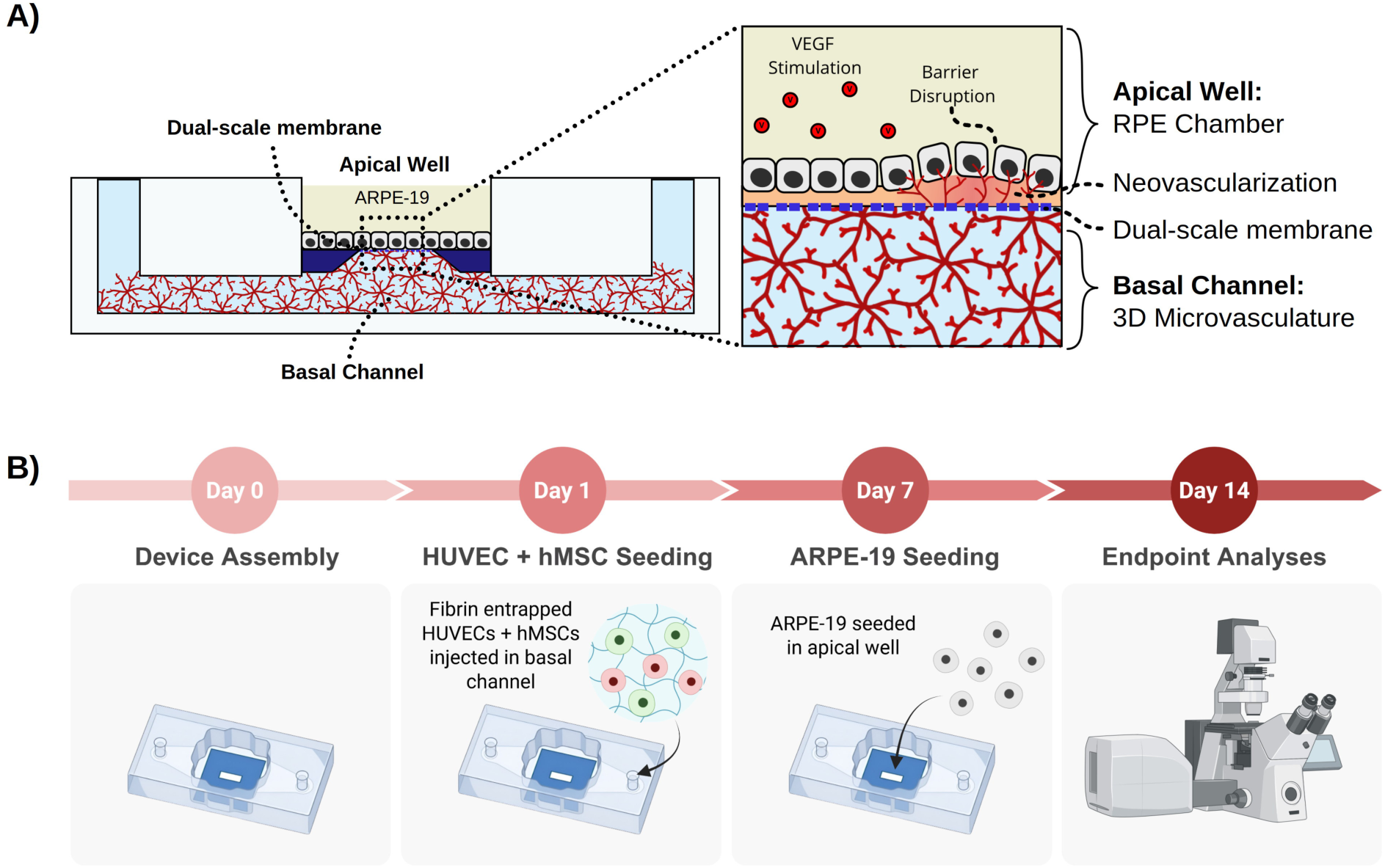
*µSiM-OBRB enables in vitro juxtaposition of 3D choriocapillaris-like microvasculature with functional ARPE-19 barrier tissues.* **(A)** Schematic of ARPE-19 barriers juxtaposed with fibrin encapsulated microvasculature at the nanomembrane interface. ARPE-19 cells form tight barriers on top of a nanomembrane. 3D choriocapillaris is formed in the basal channel, directly underneath ARPE-19 barriers. VEGF is added to the top well to induce neovascularization. **(B)** Experimental protocol for forming 3D choriocapillaris-like vasculature in µSiM-OBRB.

**Fig. 7.**
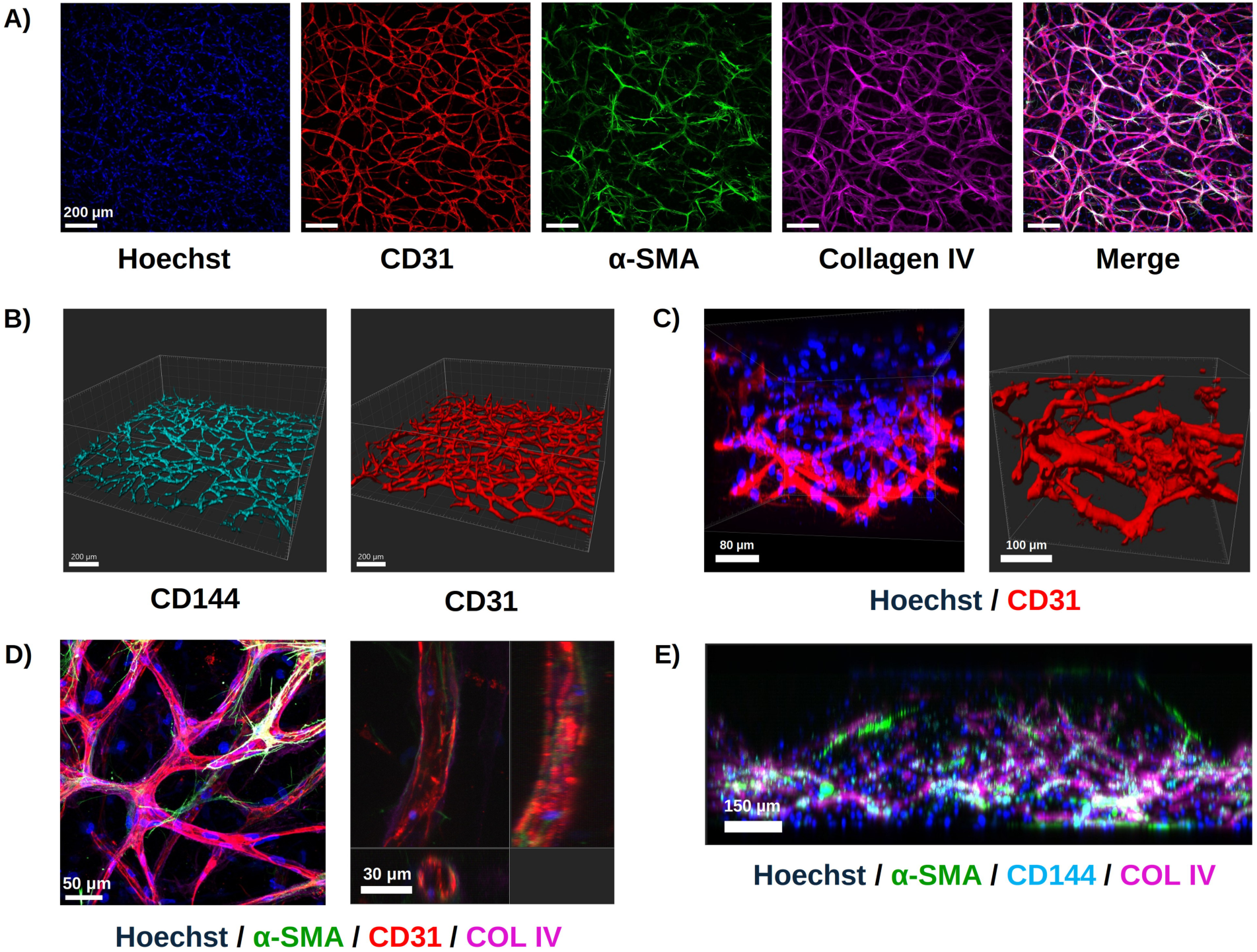
*HUVECs and hMSCs co-encapsulated in fibrin hydrogels form 3D microvascular networks in µSiM-OBRB devices.* **(A)** Max projection confocal Z-stack images depicting 3D vessel networks in µSiM-OBRB chips. Blue: Hoechst, Red: CD31, Green: α-SMA, Magenta: Collagen IV. Scale bar: 200 µm. **(B and C)** 3D views of confocal images of CD144+ or CD31+ vasculature formed in µSiM-OBRB chips. Blue: Hoechst, Cyan: CD144, Red: CD31. Scale bars: 200 µm, 80 µm, 100 µm. **(D)** Higher magnification view of max projection confocal Z-stack images and orthogonal projections depicting 3D vessel networks in µSiM-OBRB chips. α-SMA+ hMSCs form supportive pericytes, surrounding and stabilizing CD31+ vascular networks. Vascular networks deposit Collagen IV as they remodel the matrix. Blue: Hoechst, Red: CD31, Green: α-SMA, Magenta: Collagen IV. Scale bars: 50 µm, 30 µm. **(E)** Orthogonal projection of Z-stack confocal image depicting microvasculature in a µSiM-OBRB chip. Blue: Hoechst, Green: α-SMA, Cyan: CD144, Magenta: Collagen IV. Scale bar: 150 µm. All samples were fixed on day 14.

Beyond forming networks, the µSiM-OBRB microvasculature displayed several hallmarks of mature, organized vessels. hMSCs (stained with α-SMA) surrounded CD31+ endothelial cells, suggesting mural cell differentiation to support vascular stability (**Fig. 7A, D**), while endothelial networks exhibited pericellular collagen IV deposition, indicating vascular basement membrane development (**Fig. 7A, D**).

Primitive lumen formation was observed in some vessels formed in µSiM-OBRB (**Fig. 7B**). 3D µSiM-OBRB microvascular networks were formed throughout the full volume of the basal channel (**Fig. 7C–E**) (**Table S4†**).

### VEGF stimulation induces transepithelial endothelial migration, mimicking choroidal neovascularization

To mimic the RPE-choriocapillaris interface, fibrin-encapsulated HUVEC + hMSC choriocapillaris networks were cultured in the basal channel of µSiM-OBRB devices with dual-scale silicon nitride (DSSN) or microporous silicon nitride (MPSN) membranes, and ARPE-19 cells were seeded in the apical well after 7 days of microvascular development (**Fig. 8A**). RPE65+ ARPE-19 cells formed epithelial barriers in the apical chamber for an additional 7 days (**Fig. 8B**). µSiM-OBRB tri-cultures were treated with 25 ng mL^-1^ VEGF to test whether VEGF drives endothelial transepithelial migration into the apical chamber. The large 3 µm diameter pores of DSSN membranes allow for active apicobasal cellular transmigration but prevent cells from passively crossing the membrane through random migration.^52,67,68^ To track endothelial cell migration, µSiM- OBRB tri-cultures were stained for CD31. µSiM-OBRB chips with 0.5 µm pore diameter MPSN membranes that prevent transepithelial cell migration were included as a negative control.

**Fig. 8.**
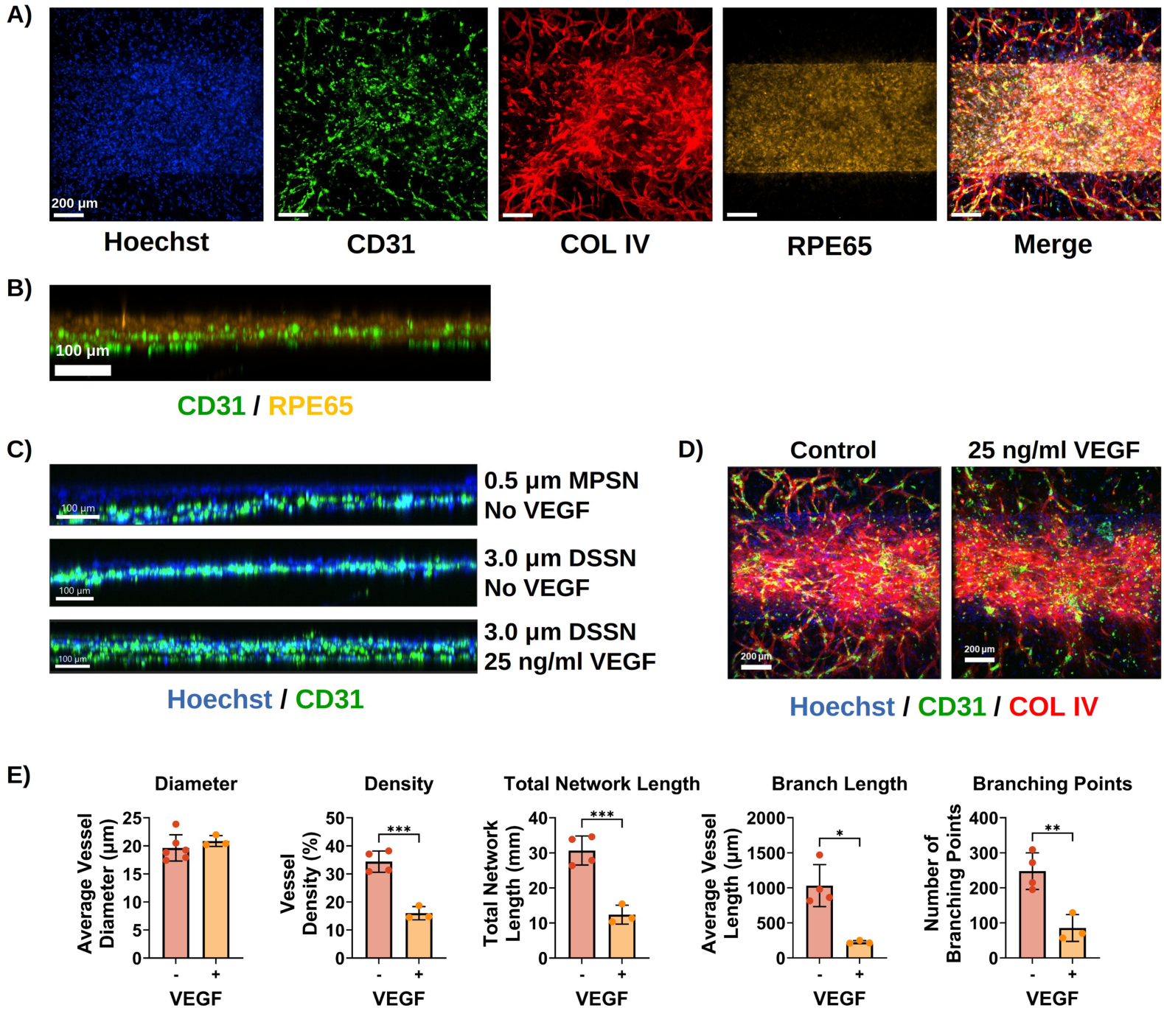
µSiM-OBRB models pathological choroidal neovascularization *in vitro.* **(A)** Z- stack maximum projection confocal images depicting RPE65+ ARPE-19 barriers juxtaposed with CD31+ and Collagen IV+ microvascular networks in the basal channel in a µSiM-OBRB device with a dual-scale membrane. ARPE-19 cells were isolated to the membrane window, where they interfaced with basal microvascular networks. Blue: Hoechst, green: CD31, red: Collagen IV, yellow: RPE65. Scale bars: 200 µm. **(B)** Orthogonal projection of a Z-stack confocal image depicting RPE65+ ARPE-19 cells above CD31+ HUVECs. **(C)** Orthogonal projections of Z-stack confocal images depicting ARPE-19 + HUVEC + hMSC tricultures in µSiM-OBRB devices with non- VEGF-stimulated MPSN, non-VEGF-stimulated DSSN, and VEGF-stimulated DSSN membranes. **(D)** Z-stack maximum projection confocal images depicting microvascular networks in VEGF stimulated and non-stimulated µSiM-OBRB chips with DSSN membranes. Blue: Hoechst, green: CD31, red: Collagen IV. In VEGF-stimulated DSSN membrane devices, CD31+ HUVECs migrated through the membrane and invaded the apical retinal space, becoming interspersed with ARPE-19 cells. **(E)** Average microvessel diameter, density, total network length, average vessel length, and number of branching points of 3D microvascular networks formed in VEGF stimulated (+) and non-stimulated (-) µSiM-OBRB devices with DSSN membranes on day 14. Mean ± SD. N=3-6. Statistics: one way ANOVA with Tukey’s post hoc comparisons test, *: p<0.05. **: p<0.005. ***: p<0.0005. All samples were fixed on day 14.

In VEGF-treated µSiM-OBRB chips (VEGF+) with DSSN membranes, CD31+ cells were interspersed among ARPE-19 cells in the apical well, indicating transepithelial HUVEC migration (**Fig. 8C**). Transepithelial HUVEC migration was not observed in non-stimulated MPSN or DSSN µSiM-OBRB tri-cultures. While the average vessel diameter in both groups was similar, VEGF+ µSiM-OBRB DSSN tri-cultures exhibited microvascular network atrophy, resulting in disconnected, incomplete vessel networks compared to the VEGF- controls (**Fig. 8D, E**). Compared to the VEGF+ group, microvascular networks in the unstimulated chips exhibited over 2-fold higher vascular density (Control: 34 ± 4%, VEGF+: 16 ± 2 % (mean ± SD)) and nearly 3-fold as many vessel branching points (Control: 248 ± 52 branching points, VEGF+: 85 ± 40 branching points (mean ± SD)). Total network length was about 2.5-fold higher in the untreated group compared to the VEGF+ chips (Control: 31 ± 4 mm, VEGF+: 12 ± 3 mm (mean ± SD)). Individual vessel branches were about 4.5-fold longer in the untreated group, indicating that VEGF stimulation reduced the networking capacity of endothelial cells (Control: 1,030 ± 300 µm, VEGF+: 225 ± 22 µm (mean ± SD)). These data indicate that VEGF stimulation of µSiM-OBRB microvascular networks caused endothelial network atrophy and endothelial invasion into the apical chamber, mimicking the pathological neovascularization of retinal degeneration.

## Discussion

The adaptation of the µSiM to a model of the outer blood-retinal barrier is a key advancement in our effort to build an in vitro tool for retinal drug development. The need for such a tool arises from two directions at once; drugs prescribed for conditions that have nothing to do with the eye reach the retina and damage it, while drugs intended for the retina cross the barrier so poorly that no tolerable systemic dose reaches the tissue, which is why the standard of care in neovascular AMD remains repeated intravitreal injection. While the compounds and the diseases change with context, the same tissue sets the limit in every case. The retinal pigment epithelium is held apart from the fenestrated choriocapillaris by Bruch’s membrane alone, and that capillary bed carries one of the highest blood flows per unit tissue weight in the body,^33–38^ so a circulating drug arrives at the epithelium having met little vascular resistance on the way. In this arrangement, the choroidal vasculature acts as the carrier, while the selectivity that determines what reaches the neural retina is imposed by the epithelium and the thin matrix beneath it. Hence, a model of this interface must place the two tissues in apposition rather than in adjacent compartments.

Adapting the µSiM to this interface is useful for multiple reasons. First, the silicon nitride nanomembrane is roughly 100 nm thick, thinner than the basal lamina it supports, so that RPE seeded on one face and endothelium seeded on the other are held in direct apposition rather than separated by a polymer slab. Models that interpose a track-etched membrane tens of micrometers thick where Bruch’s membrane belongs are interposing a structure whose barrier consequences are not well characterized. Critical among the practical benefits of the nanomembrane is that it is glass-like and the device sits on a conventional microscope, so that a monolayer can be inspected in place at any point in the culture and then fixed and stained in the device with no further handling.^46,54^ A Transwell offers neither continuous monitoring nor imaging in place, since its membrane has to be cut out and mounted for confocal imaging, introducing handling steps where tissue is easily damaged.^81^ 10 to 30 µm thick transwell membranes also scatter light, reducing resolution and ruling out imaging of juxtaposed tissues in co-culture.^82–84^ Modularity is a further advantage, and is leveraged here, since a single device body accepted nanoporous, microporous, and dual-scale membranes, a perfusable basal channel, and an injectable hydrogel compartment, so that multiple hypotheses could be tested with the same rapid assembly steps and without redesigning the system.

Our work here confirms that the µSiM supports the formation of a functional RPE barrier between a vascular and a retinal compartment, which is the most fundamental requirement of the model. We chose ARPE-19 as the RPE source for reliability and convenience during this development project, reserving the more complex and more expensive iPSC-derived RPE for the next phase of development. Under nicotinamide-supplemented medium from day 7, ARPE-19 on laminin-coated nanomembranes formed confluent monolayers with ZO-1-positive tight junctions and hexagonal cobblestone packing, with ezrin localized apically relative to nuclei and ZO-1, and with collagen IV deposited basally (**Fig. 2C–F**). Because ezrin and collagen IV segregated to opposite faces of the layer, this apicobasal sorting indicates that the cells had polarized rather than simply reached confluence, which is the state a transport measurement requires. Nicotinamide was essential to reach that state,^55^ and without it ARPE-19 stayed spindle-shaped and migratory and failed to develop cobblestone morphology, tight junctions, or apicobasal polarity (**Fig. S2†**). Notably, the nicotinamide requirement was obvious from routine inspection of the cultures rather than from an endpoint assay, because a µSiM culture can be examined directly on a microscope stage.

We also show that TEER and small-molecule permeability place the maturation of these barriers within the range of the published record. TEER in the 28-day monoculture sat above most of the values reported for ARPE-19 on microporous poly(ethylene terephthalate) or polycarbonate Transwell inserts^85–88,96–98^ (**Table S3†**) and was comparable to human donor RPE tissue,^89^ while remaining below that of the longest ARPE-19 cultures^90^ and of cultured primary RPE.^91^ Permeability behaved the same way, with our monocultures and co-barriers falling within the range reported for small-molecule transport across ARPE-19^92^ and below the values reported for microphysiological system (MPS) models of the blood-brain barrier.^46,93,94,99^ A barrier matured in weeks on a convenient cell line does not reach the values that long culture and better cell sources deliver, but the maturation we obtain is sufficient for the transport measurements that establish a working prototype of the µSiM-OBRB. Maturation was deliberately traded for a shorter culture, because our proof-of-concept work needed iteration more than maximal barrier behavior, and the use of iPSC-derived RPE is a goal for the next generation of the model. Significant among our findings is that seeding HUVECs on the basal face of the same 100 nm membrane, in direct apposition to the RPE, produced co-cultures that reached the permeability of a 28-day monoculture in half the time (**Fig. 3E**). Time to a competent barrier is the rate-limiting cost of any barrier-tissue MPS, and establishing minimum-time protocols is a critical step for our ongoing efforts to use these platforms for screening rather than for demonstration.

A primary objective for the µSiM-OBRB as a drug development tool was to validate it for bioavailability prediction. Passive transcellular transport across a tissue barrier is expected to sort small molecules by lipophilicity, and that sorting has been reported for the outer blood-retinal barrier in ARPE-19,^92,100^ in stem-cell-derived RPE,^92^ and in animals.^78,79,101,102^ To test whether the µSiM-OBRB displays this selectivity, we supplied a five-drug panel spanning more than four log units of lipophilicity (**Table 2**) through the flowing basal channel. Here, we show that uptake into the retinal chamber was low for the hydrophilic compounds and rose steadily with lipophilicity across the panel (**Fig. 4B–D**), the behavior expected of a barrier in which transport is dominated by passive transcellular movement. Importantly, the panel reached the barrier from a perfused vascular channel rather than from a stagnant donor well, a configuration no static filter assay can offer. Five compounds establish a direction rather than a predictive relationship, but the device reproduced the ordering the barrier imposes in vivo with drug arriving by the vascular route. Our work also illustrates how the platform would be used in practice, since a candidate can be run simultaneously against a reference panel, with the retinal chamber sampled and transferred to mass spectrometry, so that candidates are ranked on measured transport rather than on predicted descriptors.

Our second objective was to demonstrate the detection of drug-induced barrier injury. Retinal toxicity is usually a side effect of a drug prescribed for an indication other than a retinal one, and it is described in terms of the retinal layers that fail, most often the outer retina, rather than in terms of the barrier.^4–6^ Occasionally, however, the toxicity is directed at the barrier itself, although it is usually detected as retinal failure, because the outer retina depends on the epithelium for the ionic, fluid, and metabolic support that keeps photoreceptors viable. Hydroxychloroquine is the familiar case, accumulating in the RPE and increasing the permeability of the epithelial layer, and yet the retinopathy that ends the prescription is recognized clinically as a loss of the outer retina rather than as the barrier failure that preceded it.^7–13^ Toxicity of this kind is difficult to catch by other means, since it is invisible to a viability assay on dissociated cells and slow relative to the length of a clinical trial, and since an animal study is among the least likely settings in which to resolve it.

Here, we show that supratherapeutic digoxin produces an injury phenotype in the µSiM-OBRB (**Fig. 5**). We discovered this while examining digoxin in a preliminary transport screen, where a one-hour perfusion showed that the co-barrier reduced its transport into the retinal chamber along with that of lucifer yellow and urea (**Fig. S6†**). We then found that a supratherapeutic exposure killed cells within the barrier and opened it. Digoxin carries a documented clinical association with disturbed color vision and blindness,^60,61,95,103–105^ although that association has never been located at the RPE. Because digoxin inhibits Na+/K+-ATPase,^106^ the pump that maintains the ion gradients on which both cell viability and the transepithelial gradient depend,^107^ depolarization of the epithelium is one route that could account for the loss of barrier function we observed. Localizing Na+/K+-ATPase and scoring markers of epithelial-to-mesenchymal transition after exposure, against a non-inhibitory glycoside control would elucidate the source of the observed effects. Our focus here is the capability of the device rather than the pharmacology of digoxin.

Our third goal for the µSiM-OBRB was to display the neovascular phenotype associated with disease states such as neovascular AMD and diabetic retinopathy. In these pathologies, VEGF imbalance produces a paired lesion that is difficult to model in a single preparation, since the choriocapillaris loses density and perfusion^41,45,108^ while endothelium breaches the RPE and invades the retinal compartment.^109^ In the vascularized configuration of the device, where fibrin-encapsulated HUVECs and hMSCs formed networks with primitive lumens, mural-cell support, and matrix remodeling in the basal channel, both halves of the lesion appeared as a response to pathological VEGF stimulation: microvascular networks atrophied while CD31+ endothelial cells appeared among the ARPE-19 in the apical well. The next question is whether anti-VEGF agents, the standard of care in neovascular AMD, reverse that phenotype in a human tissue construct, which will require methods to quantify the vascular invasion.

Three limitations constrain the µSiM-OBRB as it stands, beginning with the cell source. ARPE-19 cells differ transcriptomically from native RPE and form weaker barriers than primary or stem-cell-derived cells,^31,98,110^ so the barrier values reported here should be read as a floor rather than as a ceiling, and replication in iPSC-derived RPE would establish that the transport result generalizes beyond this cell line. A second limitation is the absent neural retina, since the model carries no photoreceptors or inner retinal neurons and therefore cannot address RPE-photoreceptor interactions such as outer-segment phagocytosis. The µSiM design features an open apical well that can accept a retinal organoid^44,111–113^ in the next stage of development.

The route of administration in the toxicity experiment is a third limitation. We applied digoxin to the apical chamber, while in vivo it reaches the RPE from the choroid and is actively cleared from the retina by transporters including MDR1, BCRP, MRP2, and OAT4,^78,114–119^ and we will repeat that exposure under basal perfusion as we did in the transport screen (**Fig. S6†**). We likewise did not measure barrier function in the tri-culture, because the established µSiM TEER and permeability methods do not yet account for a hydrogel matrix in the basal channel,^46,54^ although extending the impedance module used for the monoculture measurements to that configuration would supply TEER^56^ and longitudinal confocal imaging would supply permeability.^46,120^ Nor did we verify vessel perfusability in the static configuration, where interstitial pressure gradients are the established route to it.^65,121–123^ Because the dual-scale membranes were not visible in confocal Z-stacks, we could see transepithelial invasion but not readily quantify it, a measurement that live phase imaging could supply directly.^46,49,67,120^

Future work will advance the model on each of these fronts by moving the epithelium to iPSC-derived RPE, adding a retinal organoid above it, and perfusing the vascular network below. Building the epithelium and the endothelium from a single donor would open a further line of work, since an isogenic construct would let genetic susceptibility to retinal drug toxicity be studied in a patient-specific system. Building systems that emulate human disease mechanisms and predict therapeutic response requires stepwise developments such as this one. Here, we demonstrate that the µSiM is a well-positioned MPS platform for this work, and that direct epithelial-endothelial juxtaposition across an ultrathin membrane, with drug delivered by the vascular route under flow, is an essential element of the retinal models that follow.

## Conclusions

Our report introduces the µSiM-OBRB as a platform for drug development. We illustrate its use to deliver a candidate compound by a vascular route, measure what crosses the barrier, and detect drug-induced barrier injury. We also establish that the common ARPE-19 cell line matured in the device to a polarized, junction-competent barrier, and that juxtaposing that barrier with endothelium across the nanomembrane halved the time required to reach it. Under continuous perfusion of the vascular channel, those co-barriers sorted a small-molecule panel by lipophilicity in the direction reported in vivo. Digoxin produced a loss of viability and of barrier integrity in the co- barrier, establishing that the platform detects toxicity directed at the barrier itself and precedes the retinal failure that can manifest downstream of barrier injury.

A vascularized configuration of the same device extends the platform to use with drugs targeting choroidal disease, with vascular networks in the basal channel and endothelium that invades the epithelial compartment under pathological VEGF. Perfusable 3D vasculature, engineered alternatives to fibrin, and barrier measurement in tri-culture are the developments we are pursuing next. The µSiM-OBRB is already a human-relevant platform for studying the outer blood-retinal barrier in health and in disease, and for drug development it means that a candidate can be tested against a human barrier under flow at a stage when the result can still shape the animal studies that follow.

## Author contributions

Kevin Ling: conceptualization, data curation, formal analysis, investigation, methodology, resources, software, validation, visualization, writing—original draft, writing—review & editing. Jordan Jones: data curation, investigation. Gram Hepner: data curation, investigation, software. Ahmet Gurcan: data curation, investigation, methodology, resources. Rufaro Gamariel: data curation, investigation. Andres Muriel- Torres: data curation, investigation. Meng-chun Hsu: investigation. Mehran Mansouri: investigation. Sami Farajollahi: investigation. Vinay Abhyankar: methodology, resources, supervision. Ruchira Singh: conceptualization, funding acquisition, resources, writing— review & editing. Danielle Benoit: project administration, funding acquisition, supervision, resources, conceptualization, methodology, writing—original draft, writing— review & editing. James McGrath: project administration, funding acquisition, supervision, resources, conceptualization, methodology, writing—original draft, writing— review & editing.

## Data availability

The data supporting this article, including transepithelial electrical resistance and small-molecule permeability measurements, LC-MS retinal transport data, viability and barrier-integrity measurements, and vessel-network morphometrics, have been included as part of the Supplementary Information. Raw confocal image stacks, LC-MS chromatograms, and impedance spectra are available from the corresponding author on reasonable request, and will be deposited in a public repository on acceptance with the DOI added at proof. References cited in the Supplementary Information are listed in a separate reference list within that document.

## Acknowledgements

The authors would like to thank Sonal Dalvi, Nathaniel Foley, and Leah Grego for their consulting expertise in retinal biology as well as access to laboratory resources. Additionally, the authors thank Steven George and Shirure Venktesh for their technical advice for forming 3D microvascular networks in the µSiM devices. Finally, the authors thank Bradley Smith and the Center for Advanced Research Technologies for technical assistance with the liquid-chromatography mass spectrometry experiments used to assess retinal drug transport. This work was funded by the National Institute of Health-National Eye Institute (NIH-NEI R01 EY033192, NIH-NEI R21 EY027834, NIH AR064200, NIH UH3 DE027695), the National Science Foundation (NSF EBMS2225438), and the Center for Emerging Technologies and Innovative Sciences / Bausch & Lomb, Inc. (CEIS / B&L 2430C005). Additionally, this work benefited from the support of the Metabolomics Shared Resource at the Wilmot Cancer Institute, supported in part by the University of Rochester Wilmot Cancer Institute Support Grant (P30CA272302). The content of this work is solely the responsibility of the authors and does not necessarily represent the official views of the funding organizations listed above.

