## Supplementary Information: Figures S1-S9 and Tables S1-S4 for "Design and Application of a µSiM Outer Blood-Retinal Barrier (OBRB) Model as a Drug Development Tool"

### Supplemental Figures

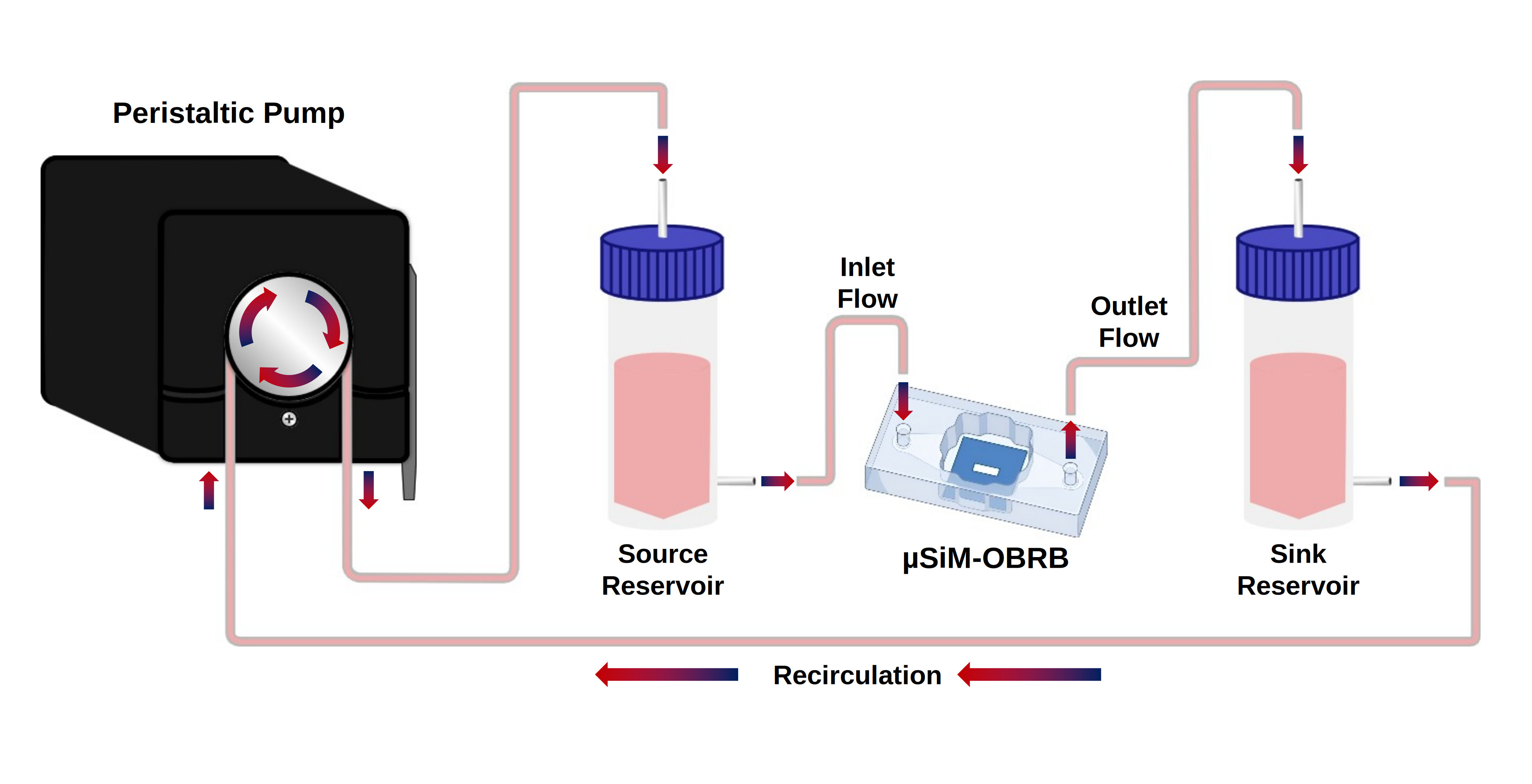

**Fig. S1** Fluidic circuit diagram for µSiM devices. A peristaltic pump set to a flow rate of 100 µL min^−1^ was used to perfuse media through the µSiM chip. 5 mL of media was added to each reservoir. Media from the source reservoir was perfused into the chip through the bottom channel, and media from the chip flowed into the sink reservoir. From the sink reservoir, media was recirculated back through the peristaltic pump and back to the source reservoir. The source reservoir also served as bubble trap.

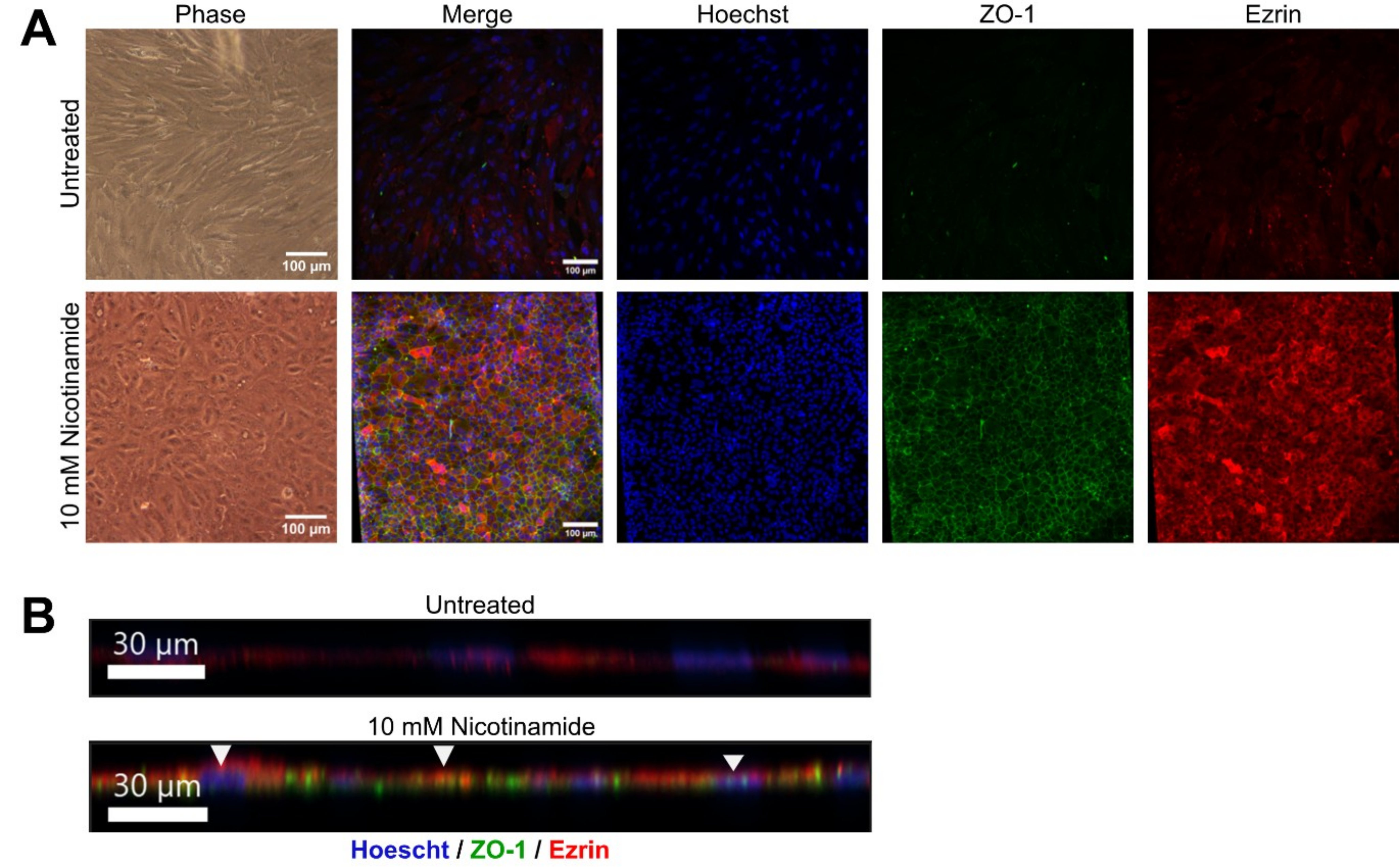

**Fig. S2** *Nicotinamide supplemented differentiation media rescues ARPE-19 differentiation, maturation, and polarization.* **(A)** Phase and confocal images of untreated or 10 mM Nicotinamide treated ARPE-19 OBRB-chip cultures. Treated ARPE-19 cultures had higher ZO-1 and Ezrin expression, and the characteristics hexagonal packing for ARPE-19 barrier tissues was clearly developed. Scale bars: 100 µm. **(B)** Nicotinamide treatment also rescues apical localization of Ezrin and the basolateral expression of ZO-1. Scale bars: 30 µm.

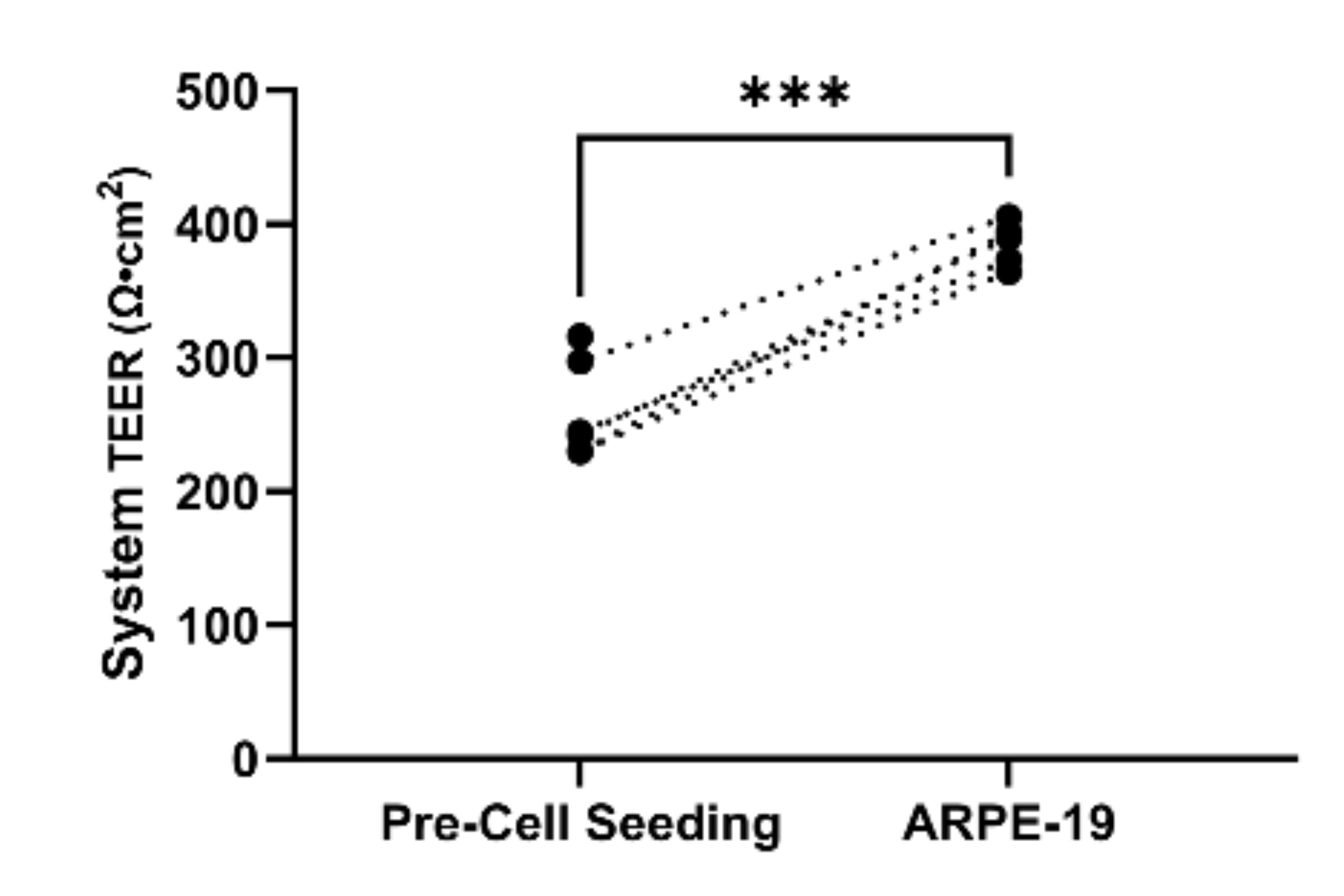

**Fig. S3** Comparison of systemic device TEER before and after ARPE-19 cell seeding and barrier formation after 28 days. Dotted lines represent paired measurements of each device tested before ARPE-19 cells were seeded and 28 days later after ARPE-19 barriers had formed on the same device. ARPE-19 barriers increased the total device systemic TEER significantly over the 4 week period.

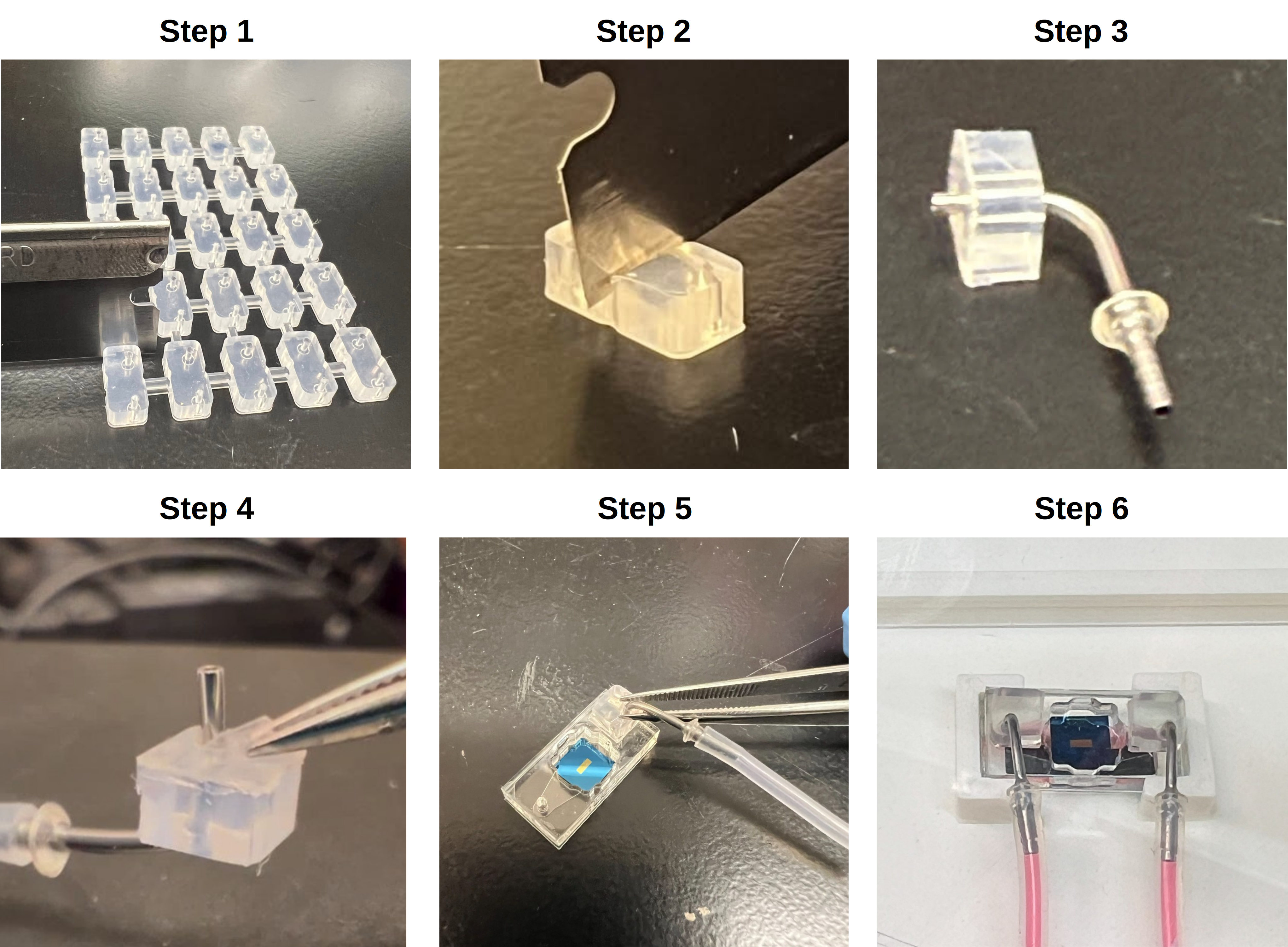

**Fig. S4** *Protocol for assembling PDMS flow bumper modules.* **Step 1:** Sterilize the PDMS flow bumper array using UV light, then use a sterile blade to cut PDMS flow bumper module from 5 x 6 array. **Step 2:** use a sterile blade to symmetrically cut the PDMS flow bumper in half perpendicular to the longest side. **Step 3:** insert a cannula into the PDMS flow bumper, allowing the tip to extend through the full flow bumper. **Step 4:** use a sharp pair of tweezers to peel the protective layer on the bottom of the PDMS flow bumper and expose the pressure sensitive adhesive. **Step 5:** Adhere the PDMS flow bumper to the top surface of a µSiM device, using the cannula tip to correctly align the flow bumper with the injection port of the basal channel. Allow to cure for 24 hours before injecting any fluids. **Step 6**: repeat Steps 1-5 and adhere a second flow bumper onto the opposite injection port.

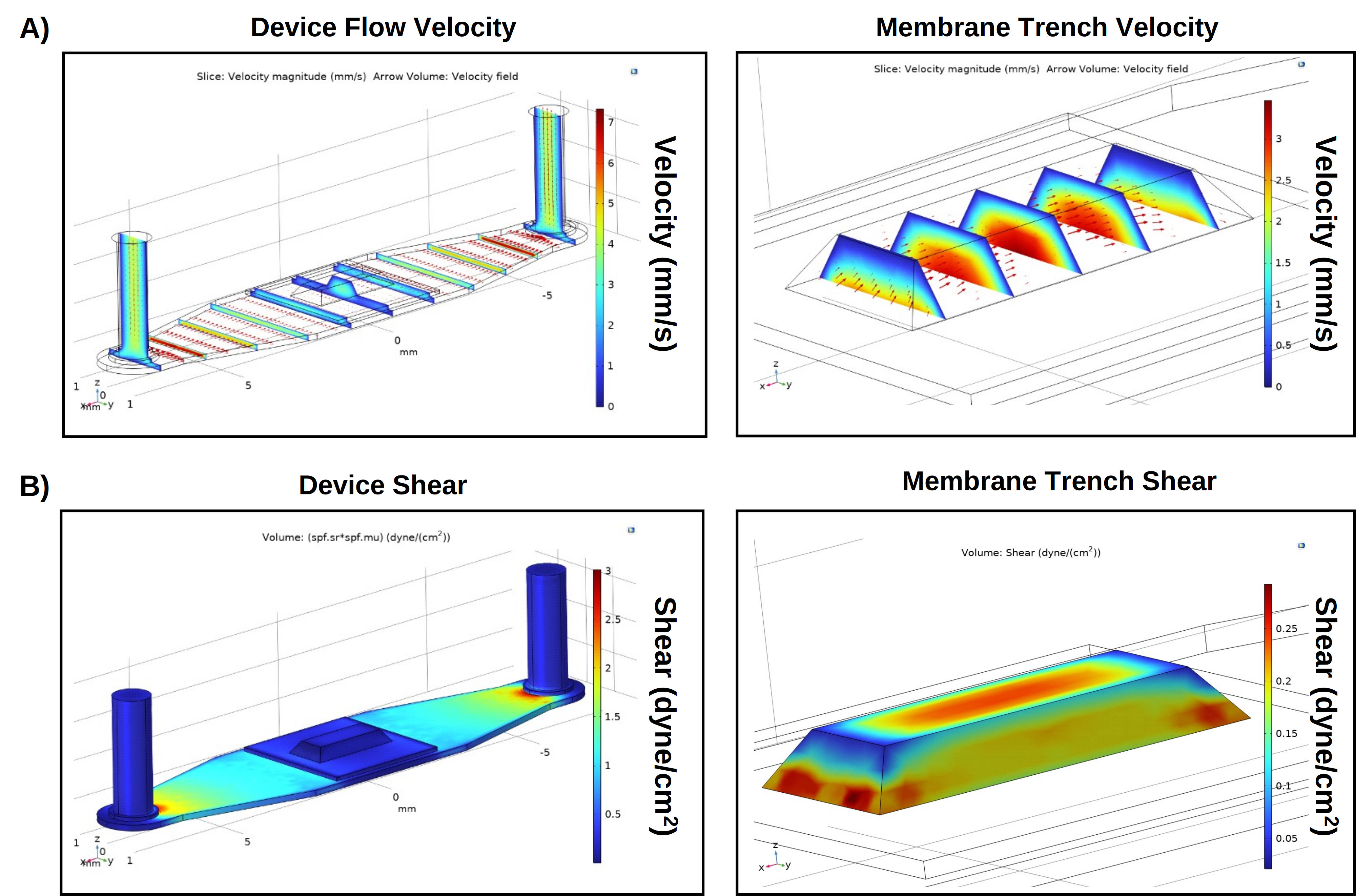

**Fig. S5** COMSOL simulations for µSiM device, assuming steady-state laminar flow. Input volumetric flow rate was set to 100 µL min^−1^. **(A)** Magnitude of flow velocity in µSiM basal channel and within the membrane trench. Volumetric arrows represent flow profiles and the velocity field. **(B)** fluidic shear in µSiM chip basal channel and within the membrane trench.

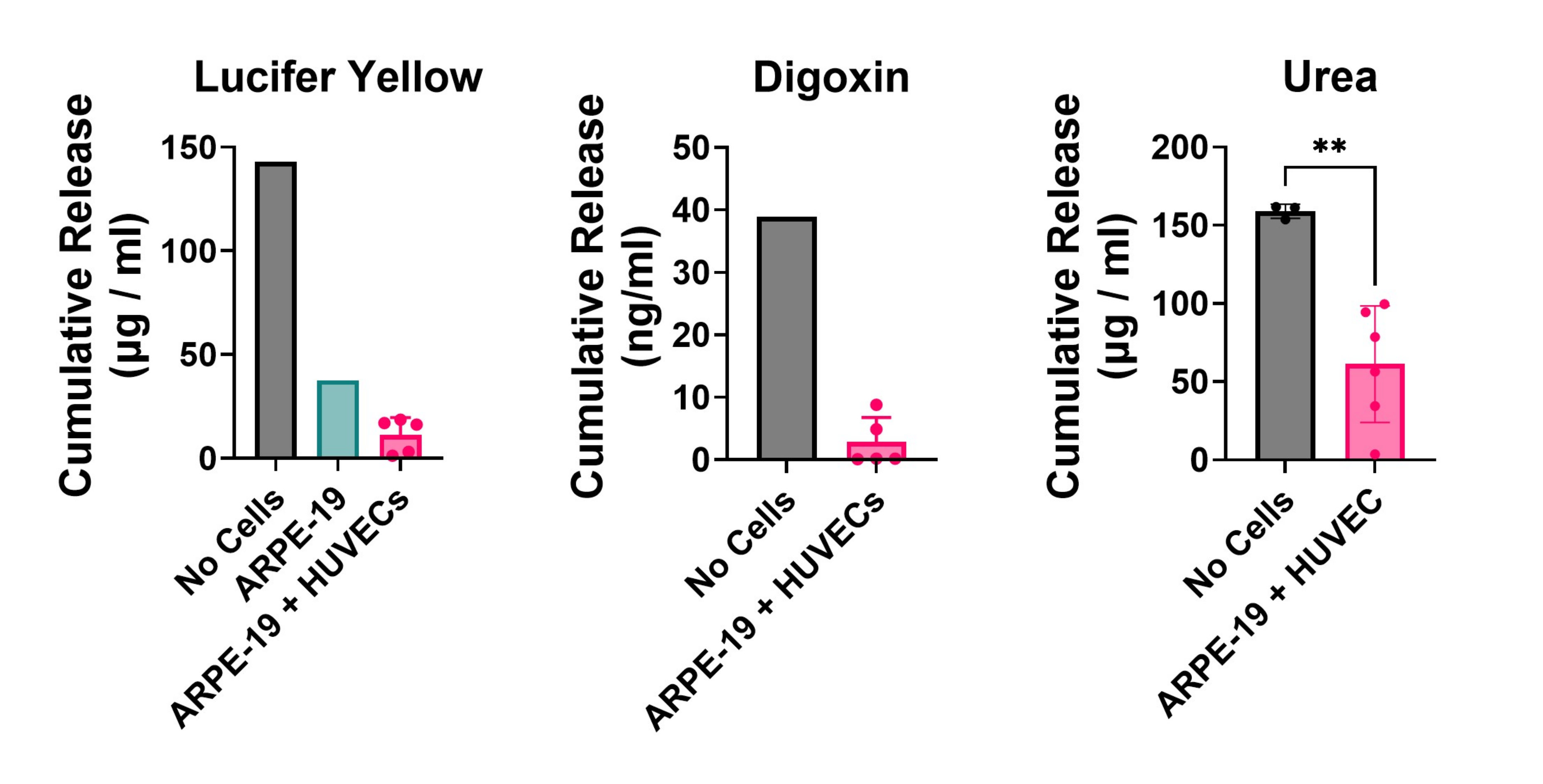

**Fig. S6** *µSiM-OBRB* *ARPE-19 + HUVEC barriers reduce retinal transport of* *small molecules in fluidic conditions.* Cumulative release of lucifer yellow, digoxin, and urea in ARPE-19 + HUVEC co-cultures, ARPE-19 monocultures, or µSiM-OBRB without cells after 100 µL min^−1^ shear flow was applied in the bottom channel for 1 hour. N=1-11. Statistics: Student’s t-test. **: p<0.005.

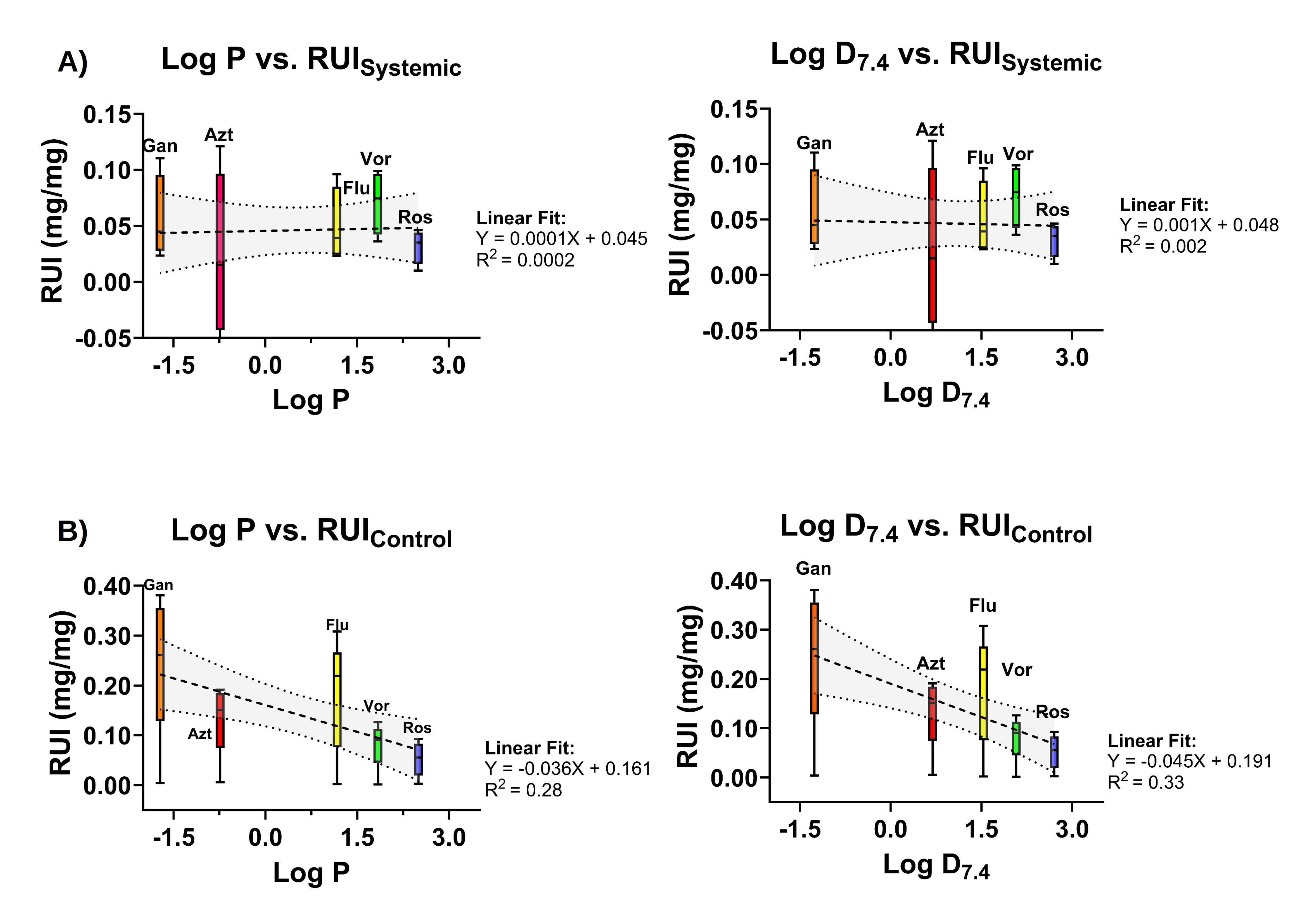

**Fig. S7** Retinal Uptake Index plotted against Log P or Log D_7.4_ for **(A)** systemic conditions (*RUI_Systemic_*) which include cell barriers cultured on µSiM devices, or **(B)** control devices (*RUI_Control_*) without cell cultures.

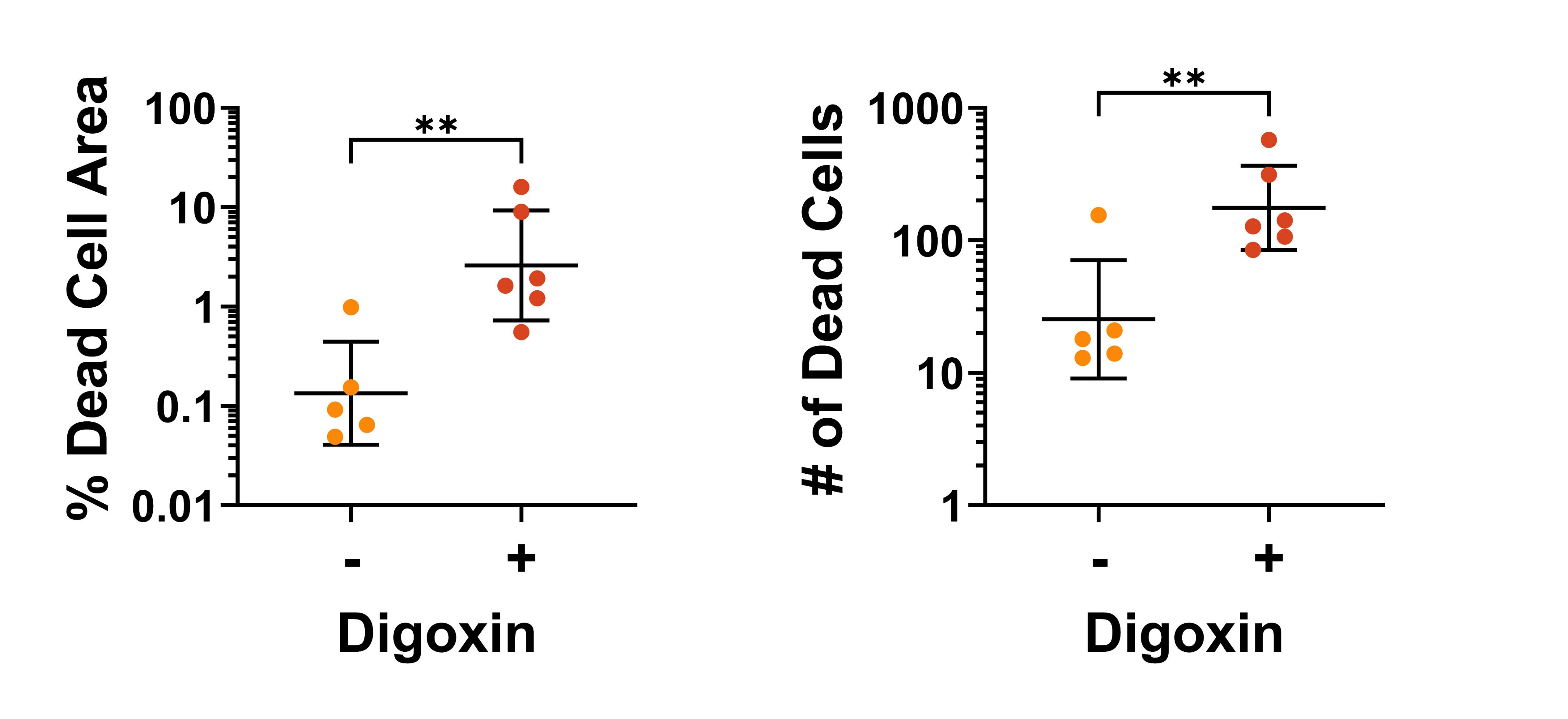

**Fig. S8** Additional quantification of LIVE/DEAD assay in digoxin treated (+) or control µSiM-OBRB ARPE-19 + HUVEC co-cultures. Left: % area of membrane surface with positive signal for the dead cell marker BOBO-3 Iodide. Right: number of distinct dead cells that were counted within the membrane region of interest. Geometric mean ± geometric SD. N=4-11. Statistics: student’s t-test. *p<0.05, **p<0.005

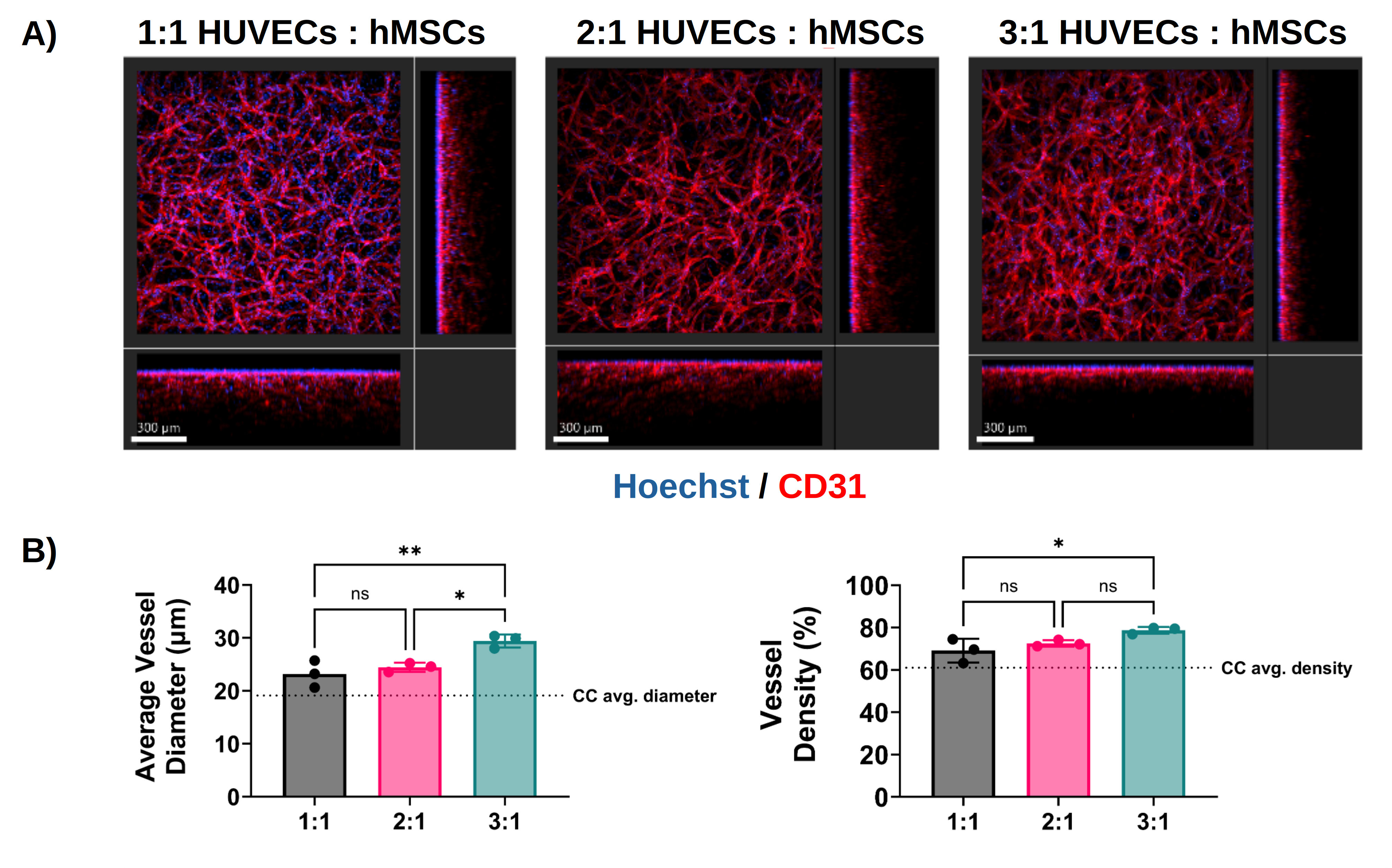

**Fig. S9** *1:1 seeding ratio of HUVECs to hMSCs produces choriocapillaris-like microvasculature in fibrin hydrogels*. (A) Confocal Z-stack max projection images of fibrin encapsulated HUVEC + hMSC microvascular networks in 96-well plates at three different cell seeding ratios. Scale bar: 200 µm. **(B)** Quantification of average vessel diameter and density of fibrin encapsulated microvascular networks at three different ratios: 1:1 HUVECs:hMSCs, 2:1 HUVECs:hMSCs, and 3:1 HUVECs:hMSCs. N=3. Statistics: one way ANOVA with Tukey’s post hoc comparisons test, *: p<0.05. **: p<0.005.

### Supplemental Tables

| **Table S1** Clinically approved drugs indicated for non-retinal conditions that have been linked to retinal toxicity. Each agent must cross the inner or outer blood-retinal barrier to reach the retina, either by permeating the intact barrier or by damaging it. Reference numbers correspond to the supplementary reference list below. | | | |
| --- | --- | --- | --- |
| **Drug or class** | **Primary indication** | **Reported retinal finding** | **References** |
| Digoxin | Heart failure, atrial fibrillation | Acute blindness, disturbed color vision, photoreceptor dysfunction, retinal degeneration | 1-5 |
| Quinine | Malaria | Retinal edema and atrophy following overdose | 6, 7 |
| Hydroxychloroquine and chloroquine | Malaria, rheumatoid arthritis, lupus | Melanin binding and retinal accumulation, increased RPE permeability, RPE cell death, bull’s-eye maculopathy; screening burden in long-term users | 8-18 |
| MEK inhibitors | Oncology | Subretinal fluid, serous retinal detachment | 19, 20 |
| FGFR inhibitors | Oncology | Serous retinal detachment | 19 |
| Phenothiazines | Psychiatric disorders | Pigmentary retinopathy and maculopathy | 19, 21 |
| Clofazimine | Leprosy, mycobacterial infection | Bull’s-eye maculopathy | 19, 21 |
| Dideoxyinosine (didanosine) | HIV infection | Peripheral retinal degeneration, reported in children | 22 |
| Deferoxamine | Iron overload chelation | Pigmentary retinopathy and maculopathy | 23 |
| Ritonavir | HIV infection | Progressive maculopathy with outer retinal disruption | 24 |
| Alkylating agents | Oncology | Maculopathy and retinal pigment epithelial change | 19, 21 |
| Denileukin diftitox | Cutaneous T-cell lymphoma | Retinopathy with visual loss | 25 |

No clinically approved treatment fully reverses toxicity-induced retinopathy; stem-cell-based regenerative approaches are in clinical trials (26-32).

| **Table S2** List of antibodies used for immunohistochemistry images, including host species, clonality, conjugates, suppliers, catalog numbers, research resource identifiers (RRID), and dilution. Primary and secondary antibodies were diluted at 1:500 v:v, and pre-conjugated antibodies were diluted at 1:200 v:v. | | | | | | | |
| --- | --- | --- | --- | --- | --- | --- | --- |
| **Target** | **Host Species** | **Clonality** | **Conjugate** | **Supplier** | **Catalog #** | **RRID** | **Dilution** |
| CD31 | Sheep | Polyclonal | N/A | R & D Systems | AF806 | AB_355617 | 1:500 |
| α-SMA | Mouse | 1A4 | N/A | ThermoFisher Scientific | 14-9760-82 | AB_2572996 | 1:500 |
| RPE65 | Rabbit | Polyclonal | N/A | ThermoFisher Scientific | PA5-78414 | AB_2736536 | 1:500 |
| Ezrin | Rabbit | Polyclonal | N/A | Cell Signaling Technology | 3145S | AB_2100309 | 1:500 |
| ZO-1 | Mouse | ZO1-1A12 | N/A | ThermoFisher Scientific | 33-9100 | AB_2533147 | 1:500 |
| CRABLP | Rabbit | Polyclonal | N/A | ThermoFisher Scientific | PA5-29759 | AB_2547233 | 1:500 |
| Mouse IgG | Goat | Polyclonal | Alexa Fluor 488 | ThermoFisher Scientific | A-11001 | AB_2534069 | 1:500 |
| Mouse IgG | Goat | Polyclonal | Alexa Fluor 568 | ThermoFisher Scientific | A-11004 | AB_2534072 | 1:500 |
| Rabbit IgG | Goat | Polyclonal | Alexa Fluor 488 | ThermoFisher Scientific | A-11008 | AB_143165 | 1:500 |
| Rabbit IgG | Goat | Polyclonal | Alexa Fluor 568 | ThermoFisher Scientific | A-11011 | AB_143157 | 1:500 |
| Rabbit IgG | Goat | Polyclonal | Alexa Fluor 647 | ThermoFisher Scientific | A32733 | AB_2633282 | 1:500 |
| CD31 | Mouse | MEM-05 | Alexa Fluor 488 | ThermoFisher Scientific | MA5-18135 | AB_2539509 | 1:200 |
| ZO-1 | Mouse | ZO1-1A12 | Alexa Fluor Plus 555 | ThermoFisher Scientific | 740002MP555 | AB_3093158 | 1:200 |
| Collagen IV | Mouse | 1042 | Alexa Fluor 647 | ThermoFisher Scientific | 51-9871-82 | AB_10853027 | 1:200 |
| VE-Cadherin | Mouse | 16B1 | Alexa Fluor 488 | ThermoFisher Scientific | 53-1449-42 | AB_10753926 | 1:200 |

| **Table S3** Reported / estimated ARPE-19 *in vitro* TEER values in µSiM-OBRB and compared to transwell cultures. N=5 for µSiM-OBRB cultures. For TEER values, mean ± SEM are shown unless otherwise noted. | | | | | |
| --- | --- | --- | --- | --- | --- |
| ***Reference*** | ***Year*** | ***Culture time*** | ***Culture substrate*** | ***Media Formulation*** | ***Reported TEER (Ω·cm²)*** |
| µSiM-OBRB | - | 4 weeks | Nanoporous silicon nitride membrane (NPSN100.C-1LZ.0) | Days 1-7: DMEM/F12 + 10% FBS  Days 7-28: DMEM/F12 + 1% FBS + 10 mM nicotinamide | 68 ± 26  (**Fig. 2G**) |
| Dunn et al., *Exp Eye Res* (35) | 1996 | 4 weeks | Transwell-COL  (0.4 µm | DMEM/F12 + 20% FBS | 50–100 |
| Ablonczy et al., *IOVS* (36) | 2011 | 2 weeks | Transwell  (0.4 µm) | DMEM/F12 + 1% FBS | 35–55 |
| Ahmado et al., *IOVS* (37) | 2011 | 3, 4, and 6 weeks (average) | Transwell  (0.4 µm) | DMEM/F12 + 1% FBS | 45.4 ± 0.8 |
|  |  |  | Transwell  (0.4 µm) | DMEM + 4.5 g/l glucose + 1% FBS | 51.8 ± 0.7 |
|  |  |  | Transwell  (0.4 µm) | DMEM + 4.5 g/l glucose + 1mM sodium pyruvate + 1% FBS | 51.3 ± 1.0 |
| Dahrouj et al., *J Pharmacol Exp Ther* | 2013 | 2 weeks | Transwell  (0.4 µm) | DMEM/F12 + 1% FBS | 40 ± 4 |
| Desjardins et al., *PLoS One* | 2016 | 2 weeks | Transwell  (0.4 µm) | DMEM/F12 + 1% FBS | 41 ± 6 |
| Samuel et al., *Mol Vis* | 2017 | 4 months | Transwell  (0.4 µm) | DMEM + 4.5 g/l glucose +1 mM sodium pyruvate + 1% FBS | 126 ± 26 (SD) |
| Hazim et al., *Exp. Eye Res.* (41) | 2019 | 6 weeks | Transwell  (0.4 µm) | MEM alpha with GlutaMAX + 1% FBS +1% N1 supplement, taurine (0.25 mg mL^−1^), hydrocortisone (20 ng mL^−1^), triiodo-thyronin (0.013 ng mL^−1^) + 10 mM nicotinamide | 40 |
| Wang et al., *Am J Transl Res* (42) | 2022 | 14–18 days | Transwell  (0.4 µm) | DMEM/F12 + 10% FBS | 32.62 ± 0.50 |
|  |  | 14–18 days | Transwell  (0.4 µm) |  | 30.61 ± 0.30 |
|  |  | 14–18 days | Transwell  (0.4 µm) |  | 29.90 ± 0.60 |
| Blenkinsop et al., *Front Cell Dev Biol* (43) | 2022 | 1 month | Transwell  (0.4 µm) | DMEM/F12 + 10% FBS + GlutaMAX + MEM non-essential amino acids solution + 10 mM nicotinamide | 171.3 ± 1.3 |
| Karakocak et al., *Eur J Pharm Sci* (44) | 2023 | 4 days | Transwell  (0.4 µm) | DMEM/F12 +10 % FBS | 54 ± 14 |
|  |  | 4 days | SiMPLI ceramic microporous membrane |  | 43 ± 6 |

| **Table S4** Reported / estimated native choriocapillaris vessel parameters compared to microvascular networks formed by HUVECs and hMSCs encapsulated in fibrin hydrogels at a 1:1 ratio in 96-well plates or µSiM-OBRB. Mean ± SD. N=6. | | | | |
| --- | --- | --- | --- | --- |
| **Property** | **Native Choriocapillaris** | **96 Well Plate** | **µSiM-OBRB** (non-stimulated) | **µSiM-OBRB** (VEGF-stimulated) |
| Diameter (µm) | 19.1 ± 0.4 µm | 23 ± 3 µm | 21 ± 3 µm | 21 ± 1 µm |
| Vessel Density (%) | 61 ± 0.7% | 69 ± 6% | 37 ± 9% | 16 ± 2% |
| Total Network Length (mm) | Not reported | - | 34 ± 10 mm | 12 ± 3 mm |
| Branch Length (µm) | Not reported | - | 550 ± 260 µm | 225 ± 22 µm |
| Number of Junctions | Not reported | - | 300 ± 170 junctions | 85 ± 40 junctions |
